# Cross-Tissue Specificity of Pediatric DNA Methylation Associated with Cumulative Family Adversity

**DOI:** 10.1101/2023.10.04.559423

**Authors:** Meingold Hiu-ming Chan, Sarah M. Merrill, Fizza Fatima, Julie L. MacIsaac, Jelena Obradović, W. Thomas Boyce, Michael S. Kobor

## Abstract

**Background:** Cumulative family adversity (cumulative FA), characterized by co-occurring stressors in a family context, may be biologically embedded through DNA methylation (DNAm) and contribute to later health outcomes.

**Materials & Methods:** We compared epigenome-wide DNAm associated with cumulative FA in buccal epithelial cells (BECs; *n*=218) and peripheral blood mononuclear cells (PBMCs; *n*=51) from 7-13-year-old children in Canada, accounting for sex, age, predicted cell-type proportion, and genetic ancestry.

**Results:** Higher levels of cumulative FA were associated with DNAm at seven sites, primarily in stress- and immune-related genes, only in PBMCs. Negative mother-child interaction contributed to this association.

**Conclusions:** The findings of this study suggested that PBMC DNAm can be used as a marker for biological embedding of cumulative FA.

## Introduction

Early-childhood adversity has been linked to long-term health and well-being in children. In particular, adverse family environments in early life could be risk factors for long-term negative health and behavioral outcomes, including poorer physical health, stress dysregulation, psychiatric disorders, and psychosocial problems. These early experiences may “get under the skin” through a number of biological pathways in a process known as biological embedding, thus influencing the course of children’s development and health [1,2]. Such biological embedding is potentially underpinned by epigenetic regulation, i.e., processes that may alter gene expression without changing the underlying DNA sequence. The best-characterized epigenetic mark in humans is DNA methylation (DNAm), specifically methylation at cytosine-guanine dinucleotides (CpGs), which is relatively stable and can be measured easily, and is therefore commonly used to study the biological embedding of early-life experience [3]. DNAm plays a significant role in cellular differentiation during embryogenesis, leading to distinct DNAm profiles across tissues and cells; therefore, DNAm associated with social experience could be dependent on the types of biological samples used in the analyses [3].

Although severe early-life adversity, e.g., childhood trauma, abuse, and poverty, has been a primary focus of human DNAm research and was shown to be robustly associated with DNA signatures [4], daily experience of more prevalent yet still stressful familial factors, especially cumulatively, may also be linked to health and development. For example, financial stress and parental psychopathology are more common adverse exposures among children (18.8% and 7.1%, respectively) than physical or sexual abuse (4%) [5,6]. In fact, recent research has shown that cumulative family adversity (cumulative FA) is associated with biomarkers of inflammation, cardio/metabolic systems, and endocrine systems [7,8], and emerging evidence suggests DNAm as a potential biological pathway underlying associations between cumulative FA and health outcomes [9,10]. Studies have begun to explore the associations of DNAm with cumulative FA in the prenatal, infant, and early childhood periods. Although these very early time points may be of interest for studying biological embedding due to its hypothesized sensitivity to environmental exposures [11], later childhood represents a developmental period of important psychosocial transitions, heightened stress, and peak years for onset of common mental health disorders [12,13]. Therefore, it is also important to understand whether and how cumulative FA during such important developmental periods is reflected in DNAm, which could have implications for later mental health outcomes. Furthermore, the roles of more direct parent-child interaction quality in the biological embedding of cumulative FA in children require more detailed investigation. The present study was performed to examine the epigenome-wide associations of cumulative FA in school-age children using two commonly collected tissues, buccal epithelial cells (BECs) and peripheral blood mononuclear cells (PBMCs), taking into consideration biological drivers of DNAm, such as tissue effects and roles of parent-child interaction quality.

### Epigenetic associations of cumulative FA

The relations of stressors in the family context, such as maternal depression, parenting stress, and financial stress, with health and developmental outcomes in children have been documented [14]. DNAm has been examined as a potential pathway for the biological embedding of these stressors in children. Many earlier studies primarily assessed the relation of *prenatal* psychosocial stress with DNAm in the offspring and found differential DNAm at candidate genes, such as *NR3C1* (specifically exon 1_F_) encoding the glucocorticoid receptor (GR) and *SLC6A4* encoding the serotonin transporter [10,15]. Nonetheless, there is intense debate regarding the biological meaning and interpretability of these findings [11,16]. There is also accumulating evidence indicating epigenetic signals of *postnatal* exposure to severe psychosocial adversity, such as childhood maltreatment or other traumatic events [17,18]. However, more prevalent albeit probably less severe types of family adversity during childhood may also be associated with DNAm differences, especially in a cumulative manner. It is important to elucidate the sensitivity of DNAm to less severe adverse conditions and illuminate the roles of more common types of cumulative FA experienced by a larger percentage of the pediatric population on children’s health and development.

In fact, epigenetic signals associated with some aspects of cumulative FA within community samples have been identified. For example, higher levels of subclinical parental psychiatric symptoms, such as depression, anxiety, or general internalizing/externalizing symptoms, were linked to differential DNAm of candidate genes in offspring, including *NR3C1* exon 1_F_, *11b-HSD2*, and *SLC6A3* [19–22]. In addition to parental psychiatric symptoms, financial stress is another aspect of cumulative FA has been linked to epigenetic signals, including differential DNAm in candidate genes and elements, such as *5HTTLPR* [23], *SLC6A4* [24], *LINE1*, *Alu*, *BDNF*, *GNF2*, and *C1S* [25]. The associations of parental stress in the prenatal period with DNAm in offspring have also been widely studied [10], and a review showed a consistent association between increased DNAm on *NR3C1* exon 1_F_ with parental stress across seven human studies [15]. However, the candidate gene approach has several shortcomings; for example, some candidate gene promoters (e.g., *NR3C1*) typically exhibited minimal variability, and examining a narrow set of candidate genes could prevent discovery of less well-documented yet still relevant genes [11]. In a genome-wide context, exposure to parental psychopathology during adolescence was shown to be associated with differential DNAm in adulthood [26] and positive epigenetic age acceleration (EAA), a DNAm-based estimator of biological aging [27]. Therefore, DNAm is associated with more prevalent and less severe forms of cumulative FA. However, these studies rarely accounted for the potential roles of the parent-child interaction quality in these processes.

There is accumulating evidence from human studies indicating relations of both positive (e.g., warmth, responsiveness, and touch) and negative (e.g., harsh parenting) maternal caregiving behaviors with DNAm in children [17]. However, many of these studies also relied on a similar set of candidate genes, such as *BDNF*, *NR3C1*, *SLC6A4*, and *OXTR*, and there has been limited research regarding specific caregiving behaviors in a genome-wide context [10,17]. Nonetheless, these studies provided some support for the linkage of certain caregiving behaviors, such as mother-child bonding and maternal touch, warmth, and harsh parenting, with differential DNAm across the genome [16,26,27]. Specifically, supportive parenting was found to moderate the positive link between adversity and EAA, suggesting that direct parent-child interaction may play a role in biological embedding [28]. Psychological studies also showed that an adverse family environment may often be associated with lower levels of supportive caregiving behaviors and poorer parent-child interaction quality [29]. Therefore, the present study was performed to elucidate the relations between cumulative FA, mother-child interaction quality, and DNAm in children, particularly in the context of cumulative FA.

Although previous research primarily assessed the independent associations of single stressors on pediatric development, children are rarely exposed to only *one* type of adverse experience, especially given the high level of co-occurrence between familial risk factors [30]. Therefore, a cumulative risk perspective is increasingly being adopted to conceptualize and examine aggregate roles of co-occurring stressors and thus provide a more comprehensive representation of children’s adverse experiences in the family context [30–32]. Using this cumulative environmental risk approach, both cross-sectional and longitudinal associations have been identified with a wide range of negative child health and behavioral outcomes, including internalizing and externalizing problems [30], neurocognitive development [33,34], and psychosocial adjustment [35]. These cumulative risks were shown to be biologically embedded in children, as they were associated with biomarkers of inflammation (e.g., C-reactive protein) and stress reactivity (e.g., cortisol reactivity, autonomic nervous system) [8,30,36–39].

To our knowledge, there have been only two previous epigenome-wide association studies (EWASs) regarding the relations of multiple types of family psychosocial adversity (including maternal depression, financial stress, and parenting stress) with differential DNAm in children [31,32]. Both studies were conducted in North America with racial and socioeconomic contexts similar to the present study—primarily White and homogenous, middle-class family income level. DNAm data in BECs were collected in these studies at ages 9-11 and 15, respectively, and both provided some evidence of epigenome-wide signals related to cumulative FA, with stronger signals in genes related to growth, brain development, and immunological responses, using DNAm arrays assessing approximately 270,000 [32] and 450,000 CpG sites [31]. However, a recent study found no significant association between cumulative psychosocial stress (including maternal mental health and parental relationship problems) and EAA [40]. Given the paucity of studies regarding genome-wide associations of DNAm with cumulative FA, further research using similar standardized measures of the family environment is needed to corroborate previous findings [17]. Therefore, the present study was performed to examine the biological embedding of cumulative FA using similar measures while considering important biological factors associated with DNAm, such as tissue specificity.

### Cross-tissue investigation of DNAm

As mentioned above, the only two EWASs on cumulative FA reported to date measured DNAm in buccal samples [31,32]. Similarly, the majority of epigenetic studies on early-life psychosocial adversity also used BECs [20,26,41–43]. However, given its role in cellular differentiation, DNAm profiles are distinct across tissues and cells. Therefore, tissue and cell types are major contributors of DNAm variations, and findings regarding DNAm could be specific to the tissue in which they are examined [11]. Furthermore, tissue-specific genetic influences can occur through methylation quantitative trait loci (mQTLs), genetic variants strongly associated with nearby CpGs. Although most mQTLs are consistent across tissues [44], some can show specificity and may drive differential DNAm between tissues [45]. Therefore, it is unclear whether the associations of DNAm with cumulative FA identified in previous studies were specific to BEC DNAm or shared across tissues, which could have different biological implications.

Buccal and blood samples represent two readily accessible tissue candidates commonly used in human cohorts. While human epigenetic studies are most commonly performed using blood [46,47], buccal samples can be obtained less invasively from cheek swabs and are used more frequently in pediatric cohorts [4]. Buccal samples contain two main cell types, i.e., BECs and neutrophils, whereas the cell-type composition of blood depends on the sample type. Interindividual differences in these cell-type proportions can contribute to DNAm variation and must be accounted for in DNAm analyses.

These two tissue types are both of interest and appropriate for studying biological embedding of social experience, albeit for different reasons. BECs originate from a similar germ layer to brain cells during early development and can serve as a surrogate tissue for the brain, so are of interest due to their relevance to human brain function and psychological symptoms, especially in children [47,48]. On the other hand, blood samples, especially PBMCs consisting of neutrophils, monocytes, and lymphocytes, are of interest for analysis of immune function and inflammation. There is ample evidence for links between stress and immune system outcomes, including inflammatory processes and biomarkers [49]. Therefore, blood samples are excellent candidates for analysis of such biomarkers related to stress exposure [11].

DNAm in both tissues was found to be positively associated with brain DNAm, with slightly higher correlations for buccal than blood DNAm in one study (i.e., *r*=0.88 and 0.87, respectively) [50] and vice versa in another (i.e., *r*=0.85 and 0.86, respectively) [51]. Therefore, the two tissues may be appropriate for studying biological embedding of cumulative FA in the contexts of stress and health. However, while both tissues may be appropriate, these samples have meaningful biological differences in their origin and function. A recent comprehensive cross-tissue comparison of DNAm in matched pediatric BECs and PBMCs using the same cohort as the present study, the Gene Expression Collaborative for Kids Only (GECKO) [45], indicated marked interindividual DNAm variability both within and across BECs and PBMCs. These findings highlighted the similarities and differences in DNAm profiles across matched tissues, further bolstering the importance of careful tissue selection in DNAm studies.

Nonetheless, the questions of which tissue type should be used to investigate the associations of DNAm with early-life adversity and its potential concordance across tissue types remain open due to the paucity of cross-tissue comparisons in social epigenetic studies. Although there was some recognition that PBMC samples may better capture DNAm variations associated with the social environment, at least within the hypothesis of chronic stress [31,47], there has been limited research on cumulative FA using pediatric blood samples. Therefore, further investigations are required to examine its appropriateness.

### The present study

The present study took a cumulative risk approach to examine the biological embedding of cumulative FA through DNAm in two tissues commonly obtained in pediatric DNAm research, i.e., BECs and PBMCs. Specifically, we investigated the epigenome-wide associations of cumulative FA with DNAm in BECs and PBMCs in 218 and 51 school-age children, respectively, in the Canadian GECKO cohort [45]. The inclusion of 50 matched tissue samples also provided a unique opportunity to examine tissue specificity and cross-tissue concordance of DNAm differences in relation to cumulative FA. We also assessed the roles played by mother-child interaction quality in explaining the association between cumulative FA and DNAm in children.

First, we hypothesized that DNAm levels, especially in genes involved in stress-related pathways, may be associated with cumulative FA. Second, given the strong tissue specificity of DNAm based on both biological function and as demonstrated in previous research, we expected to find unique associations of DNAm with cumulative FA depending on tissue type [11]. Finally, we hypothesized that mother-child interaction quality, especially negative interactions given the context of adversity, would explain some of the observed associations between DNAm and cumulative FA.

## Methods

### Participants

This study included 219 mother-child dyads representing a subsample of the GECKO cohort [45], a Canadian cohort of 402 families with 6-11-year-old children designed to examine how social environment was related to child development and gene regulation. Mothers were recruited from Vancouver, British Columbia, and provided informed consent for collection of survey data for themselves as well as buccal DNA samples from their children (age range 6-11 years, *n*=376) and/or blood DNA samples in a follow-up study (age range 7-13 years; 0.5-2.3 years after buccal sample collection, *n*=92). This DNAm substudy was approved by the Clinical Research Ethics Board (CREB) at the University of British Columbia.

The analyses were conducted based on a subsample of 219 children with both maternal survey and DNAm data from BECs (*n*=218; 47.2% female) and/or PBMCs extracted from blood samples (*n*=51; 43.1% female), with both BEC and PBMC DNAm available from 50 children (44.2% female). The ethnic identities of mothers and children were reported by the mothers using a multiple-choice item with 25 possible options, including “multiracial/multiethnic” and “other,” which were then collapsed into five groups: White, Black, Asian, First Nation, multiracial, and other. Mothers primarily self-reported as White (48.2% and 64.7% for buccal and blood samples, respectively), married (75.2 %), and college-educated (87.2 %) (full demographic information is presented in Table 1). In this sample, 52.8% of children were classified as White according to maternal reports. Children were aged 6.98–12.97 years (mean=9.38, SD=0.97) and 7.00–12.94 years (mean=10.8, SD=1.14) at the time of buccal and blood sample collection, respectively.

**Table 1.**
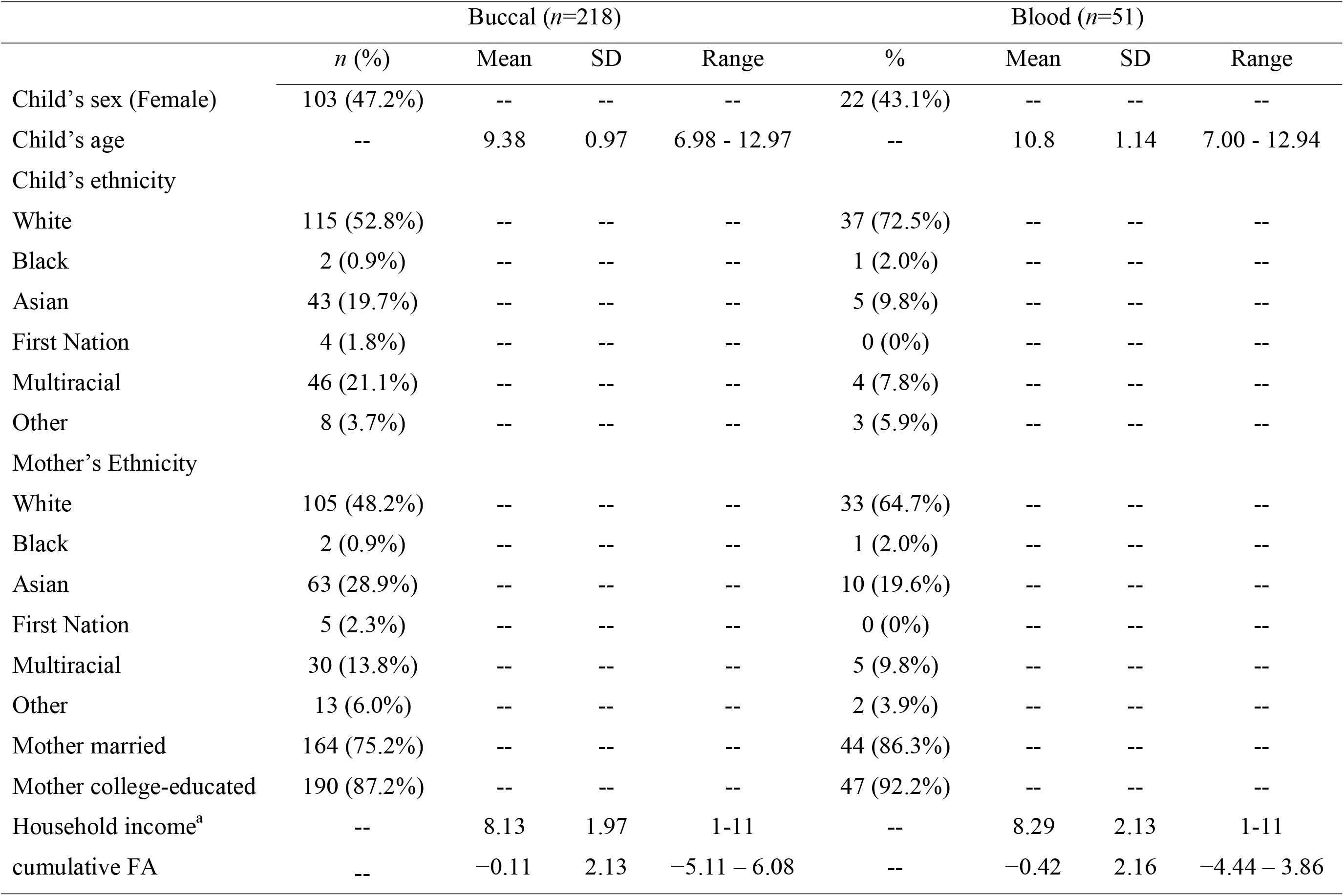
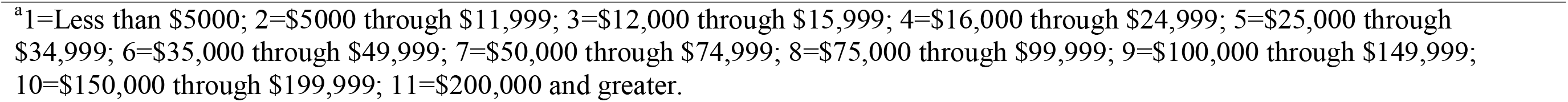
Descriptive Statistics.

### Measures

#### Cumulative FA

Measures of cumulative FA were obtained from mothers through self-reported surveys based on a composite of three family stress measures, i.e., maternal depressive symptoms, parenting stress, and financial stress, which were also used as indicators of cumulative FA in previous pediatric DNAm studies [31,32]. Maternal depressive symptoms were assessed by the Center for Epidemiologic Studies Depression (CES-D) Scale [52], which showed good reliability with these data (Cronbach’s alpha=0.83). Maternal parenting stress was measured by the Parenting Role Overload scale, which included five items regarding the amount of stress the mother experiences on a 5-point scale (e.g., “How often do things you do add up to being too much?”), and the average score was then calculated [32]. This scale also showed good reliability (Cronbach’s alpha=0.72). Maternal financial stress was assessed based on the average of four items (e.g., “How often have you thought about any current or future money problems?”) on a 5-point scale with good reliability (Cronbach’s alpha=0.80). The scores from these three scales were significantly positively correlated (*r*=0.22-0.36, *p*<.001). A composite score of cumulative FA was created by summing the standardized scores of the three scales.

#### Mother-child interaction quality

Mother-child interaction quality was assessed through both survey reports and interviews with mothers. To leverage the rich information from all parenting-relevant scales, principal component analysis (PCA) was conducted to reduce the dimensions. Measures included two surveys, the Parent Feelings Questionnaire (PFQ) [53] and Parent-Child Conflict Tactics Scale (CTSPC) [54], along with the modified Five-Minute Speech Sample (FMSS) interview [55,56].

The PFQ included 34 items eliciting responses regarding parents’ feelings about their relationship with their children. Specifically, the *negative parent-child relationship* subscale was created by averaging the mother’s rating of 13 negative statements on a 5-point Likert scale (e.g., “My child and I fight or argue more than I would like to”). Two additional subscales for parental emotions related to their children, *positive* and *negative emotions*, were created by averaging the mother’s rating on a 10-point frequency scale for 10 positive (e.g., happy) and 10 negative (e.g., angry) emotions, respectively. The reliability was good for all scales (Cronbach’s alpha=0.638-0.907).

The CTSPC assessed parental behaviors during conflict with their children. Mothers were asked to indicate how often they used each behavior on a 7-point scale (0=never; 6=more than 20 times in the past year). Ratings >2 were later recoded to represent chronicity, with 3-5, 6-10, 11-20, and >20 times in the past year coded as 4, 8, 15, and 20, respectively. Two subscales from the CTSPC, *nonviolent discipline* (Cronbach’s alpha=0.752) and *psychological aggression* (Cronbach’s alpha=0.526), were used in the analysis.

In the modified FMSS, mothers were asked to describe their children in the first 5-minute section of the interview, and then describe their relationship with their children in the second 5-minute section. Speech samples were recorded and transcribed verbatim for later coding by three trained raters. Six variables were coded from the speech samples, among which *positive comments*, *negative comments*, *general warmth*, and *general negativity* were used in the analysis. The interrater reliability was good (agreement=0.68-0.96).

PCA with oblique rotation was conducted using nine variables related to parenting behaviors. The scree plot suggested a 2-factor solution (Figure S1). The first two PCs explained 54.13% of the total variance. The first factor, *negative mother-child interaction quality*, accounted for 35.8% of the variance, with *negative parent-child relationship, parental negative emotion, nonviolent discipline, psychological aggression, general negativity, and negative comments* loaded onto the factor (Table 2). The second factor, *positive mother-child interaction quality*, accounted for 18.3% of the variance, with positive parent-child interaction quality, positive emotions, positive comments, and general warmth loaded onto the factor (Table 2). Scores of the two PCs were extracted for downstream analysis.

**Table 2.** Factor Loadings for Negative and Positive Mother-Child Interaction PCA Factors.

|  | **Negative interaction**<br>**PCA factor** | **Positive interaction**<br>**PCA factor** |
| --- | --- | --- |
| **PFQ Negative parent-child**<br>**relationship** | 0.731 | −0.290 |
| **PFQ Negative emotion** | 0.746 | −0.172 |
| **CTSPC Nonviolent discipline** | 0.806 | −0.182 |
| **CTSPC Psychological aggression** | 0.771 | −0.320 |
| **FMSS General negativity** | 0.459 | −0.812 |
| **FMSS Negative statements** | 0.421 | −0.747 |
| **PFQ Positive emotion** | −0.315 | 0.372 |
| **FMSS Positive comments** | −0.137 | 0.824 |
| **FMSS General warmth** | 0.255 | 0.516 |

*Note:* Only loadings >0.30 are shown in the table. PFQ, Parent Feelings Questionnaire; CTSPC, Parent-Child Conflict Tactics Scale; FMSS, Five-Minute Speech Sample (FMSS) interview.

### DNAm and genetic data

All procedures were approved by the institutional review board at the University of British Columbia and were described previously [45]. Full details of sample collection, extraction, DNAm and genotype profiling, and data preprocessing are presented in the Supplementary Material. Briefly, buccal samples were obtained from children using Isohelix Buccal Swabs (Cell Projects Ltd., Kent, UK) and blood samples were collected at a follow-up visit 8 months to 2.06 years later. PBMCs were isolated from the blood samples. BECs and PBMCs were applied to the Illumina Infinium HumanMethylation450K BeadChip DNAm array. Saliva samples were also collected from children to extract DNA for genome-wide single nucleotide polymorphism (SNP) profiling using the Illumina Infinium PsychChip BeadChip genotyping array. The SNP data were subjected to multidimensional scaling (MDS), and the first three MDS variables were included in subsequent analyses to account for genetic ancestry.

All tissue samples (376 BECs and 92 PBMCs, in addition to 55 dried blood spots not used in the present study) were preprocessed together to reduce the impact of batch effects. After preprocessing and removal of duplicate samples from the same individuals (*n*=10), 218 BECs and 51 PBMCs (all with genotype data), of which 50 were matched from the same individuals, and 485,577 probes in each sample were used for subsequent analyses. To reduce the testing space and address the equal significance likelihood assumption of the Benjamini-Hochberg (BH) False Discovery Rate (FDR) control method [57,58], only 270,133 and 204,753 variable probes were included in subsequent analyses for BECs and PBMCs, respectively.

## Data analysis

### Computational cell-type proportion estimation

To account for cell-type heterogeneity, a well-established driver of interindividual DNAm variation (Jones et al., 2018), we used DNAm-based methods to computationally estimate cell-type proportions in BECs and PBMCs [59]. Epithelial cell proportion was included as a covariate in subsequent analyses of BECs. For PBMCs, three PCs accounting for 92.57% of the overall variation of cell-type proportions were included as covariates in DNAm models.

### Change in DNA methylation (***Δβ***) calculation

The change in DNAm level was quantified by the change in DNAm value, or beta value (Δβ) [11]. In our analyses, Δβ was calculated by extracting the beta coefficient of cumulative FA from the robust linear regression models described above, representing the change on the y-axis (β value) divided by the x-axis (cumulative FA) while controlling for other covariates, and then multiplying the coefficient with the range of cumulative FA values between the 5^th^ and 95^th^ percentiles to reduce the effects of outliers. The resulting Δβ value represented the change in β value at each CpG associated with cumulative FA while adjusting for other covariates.

### EWAS with robust linear regression for cumulative FA

Child characteristic covariates known to be significantly associated with DNAm levels, including sex, age, genetic ancestry, and cell type, were included in the model [11]. In analyses of BECs, cell-type proportion was represented by epithelial cell proportion, while the first three cell-type PCs were included as covariates for PBMCs.

The robust linear regression model used for the site-by-site EWAS and candidate analysis was:

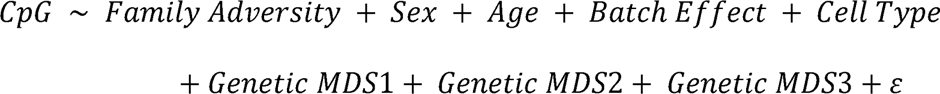

For *a priori* candidate analysis, we identified a list of biologically relevant candidate genes reported to be related to early-life psychosocial adversity and stress response: *NR3C1*, *FKBP5*, *SLC6A4*, *MAOA*, *GAD1*, *SLC6A3*, *COMT*, *OXTR*, *BDNF*, and *ADCYAP1R1* [60,61]. These genes were then used to search for 20 additional target genes sharing biological pathways but studied less frequently in previous candidate analyses using the web-based gene network and functional annotation tool GeneMANIA (http://genemania.org/) with max resultant genes set to 20 and Gene Ontology Biological Process-based weighting. A total of 30 target genes resulting in 366 and 323 variable probes for BECs and PBMCs, respectively, were included in candidate analyses (Tables S1, S2).

All variable probes were included for the exploratory EWAS (BECs: 270,133; PBMCs: 204,753). Given the large number of tests, we used the BH FDR control method for multiple-test correction [58]. A medium-confidence statistical threshold was set at FDR<0.20 corresponding to approximately *p*<2.00e−04 and *<*2.00e−05 in our candidate analysis and EWAS, respectively, to determine significant associations between DNAm at each site with cumulative FA. We also set a technical threshold of |Δβ>.03|, as it was likely to be greater than technical noise given the observations in previous studies using pediatric samples, the same DNAm array, and similar preprocessing pipelines [62,63]. We used BACON, a Bayesian method, to check for bias and inflation in our EWAS models [64], and showed no inflation of the two EWAS models in our analyses (estimated inflations of 0.95 and 0.96 for BECs and PBMCs, respectively).

### Characterization of identified differentially methylated sites associated with cumulative FA

CpGs found to be significantly associated with cumulative FA were annotated to genes, genomic regions, and chromosomal and map locations based on UCSC Genome Browser (GRCH37/hg19) [65]. We explored the biological functions and gene network interactions of the annotated genes using the GeneCards (https://www.genecards.org/) database, EWAS Atlas (https://ngdc.cncb.ac.cn/ewas), and GeneMANIA (http://genemania.org/).

#### Cross-tissue concordance & specificity

DNAm profiles are highly tissue-specific even within the same individual [11,45]. We examined the concordance of DNAm levels at the identified CpGs across BECs and PBMCs from the same individuals by Spearman’s correlation analysis. In addition, we examined whether the CpGs identified in one tissue were also differentially methylated in the other when restricted to a smaller testing space with lower multiple-testing burden. Cross-tissue information of identified CpGs was extracted from a previous study involving thorough cross-tissue comparison in the same cohort [45]. Furthermore, we assessed the blood-buccal correlation using the Iowa Methylation Array Graphing Experiment for Comparison of Peripheral Tissue and Gray matter (IMAGE-CpG) database (http://han-lab.org/methylation/default/imageCpG) [51].

To characterize the identified sites in the brain, we also examined their variability in brain and correlations between DNAm levels in blood and brain using BECon (Blood-Brain Epigenetic Concordance) (https://redgar598.shinyapps.io/BECon/) [66] and covering BA 7 (superior parietal lobule), BA 10 (frontal cortex), and BA 20 (temporal cortex) and IMAGE-CpG [51] covering the amygdala, hippocampus, and temporal cortex. The average correlations across brain regions are reported.

#### mQTL analysis

As genetic variations at some SNPs known as mQTLs are known to be associated with DNAm levels of nearby CpGs [44,45], we investigated their potential contributions to DNAm of the identified CpGs. Given the small sample size and as mQTLs are most commonly lie within 5-10 kb of a given CpG [44], we only examined variable SNPs within a 5-kb window from the identified CpGs (Table S3). The SNP genotypes were coded numerically as 1, 2, or 3 for ANOVA to examine whether DNAm level of each CpG differed across genotypes. For sites determined to be mQTLs, we reported the Δβs across SNP allele variants. In addition, we checked whether any of the CpGs were reported as mQTLs in mQTL Database (http://www.mqtldb.org/) (for PBMC CpGs) and whether they had bi- or trimodal distributions, usually signifying the effects of genetic variants, using the *nmode* function in the R package `ENmix` [67,68]. We also further examined the contributions of these SNPs to the associations between significant CpGs (where DNAm level differed across genotypes) and cumulative FA.

#### Contribution of maternal parenting to DNA methylation

We conducted contribution analysis to examine the potential contributions of maternal parenting to DNAm levels at the identified CpGs, as in previous DNAm studies [62]. The percentage contributions for negative and positive maternal parenting factors (described above) were calculated as:

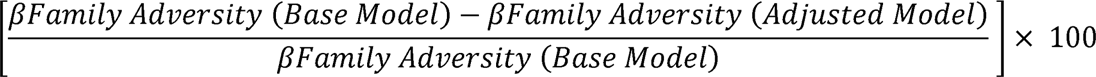

Here, the base model was the linear regression model used for candidate analysis or EWAS discovery, while the adjusted model was the base model after including a maternal parenting factor as a covariate. This was conducted for each CpG of interest identified in candidate analysis and EWAS. Two separate sets of analyses were conducted for negative and positive maternal parenting factors to examine their individual roles. All data analysis steps were performed in R version 4.2.0 and Bioconductor version 3.15.0.

#### Data and analysis code availability statement

The data for buccal samples have been deposited in NCBI Gene Expression Omnibus (GEO) (accession number GSE137903) [69]. The analysis scripts (R Markdown) used in the analyses and supporting the findings are available from the first author upon reasonable request.

## Results

### One and two candidate CpGs were associated with cumulative FA in BECs and PBMCs, respectively

A series of linear regression models of cumulative FA on DNAm were run using 366 and 323 variable probes for BECs and PBMCs, respectively (Tables S1, S2). Child’s sex, age, cell-type proportions (epithelial cell proportion in BECs and 3 PCs in PMBC), three PCs of genetic ancestry, and batch effect were included in the model. After BH correction, one CpG (cg22891314) in BECs passed the statistical threshold of FDR<0.02 (*p*<2.00e−04) but not the biological threshold of |Δβ>.03| (Figure 1A). This site was annotated to *SLC6A3* (Table 3). Two CpGs (cg00052684 and cg22659953) in PBMCs passed both the statistical and technical thresholds, operationally defined as medium-confidence site associations [32,62] with cumulative FA in our sample (Figure 1B,C). These sites were annotated to *FKBP5* and *SLC6A3* (Table 3). In GeneMANIA, 4 of 10 genes output from *FKBP5* were involved in inflammatory function and another 3 were related to the GR pathway, while 5 of 6 genes output from *SLC6A3* were associated with neurological function (Table S7).

**Figure 1.**
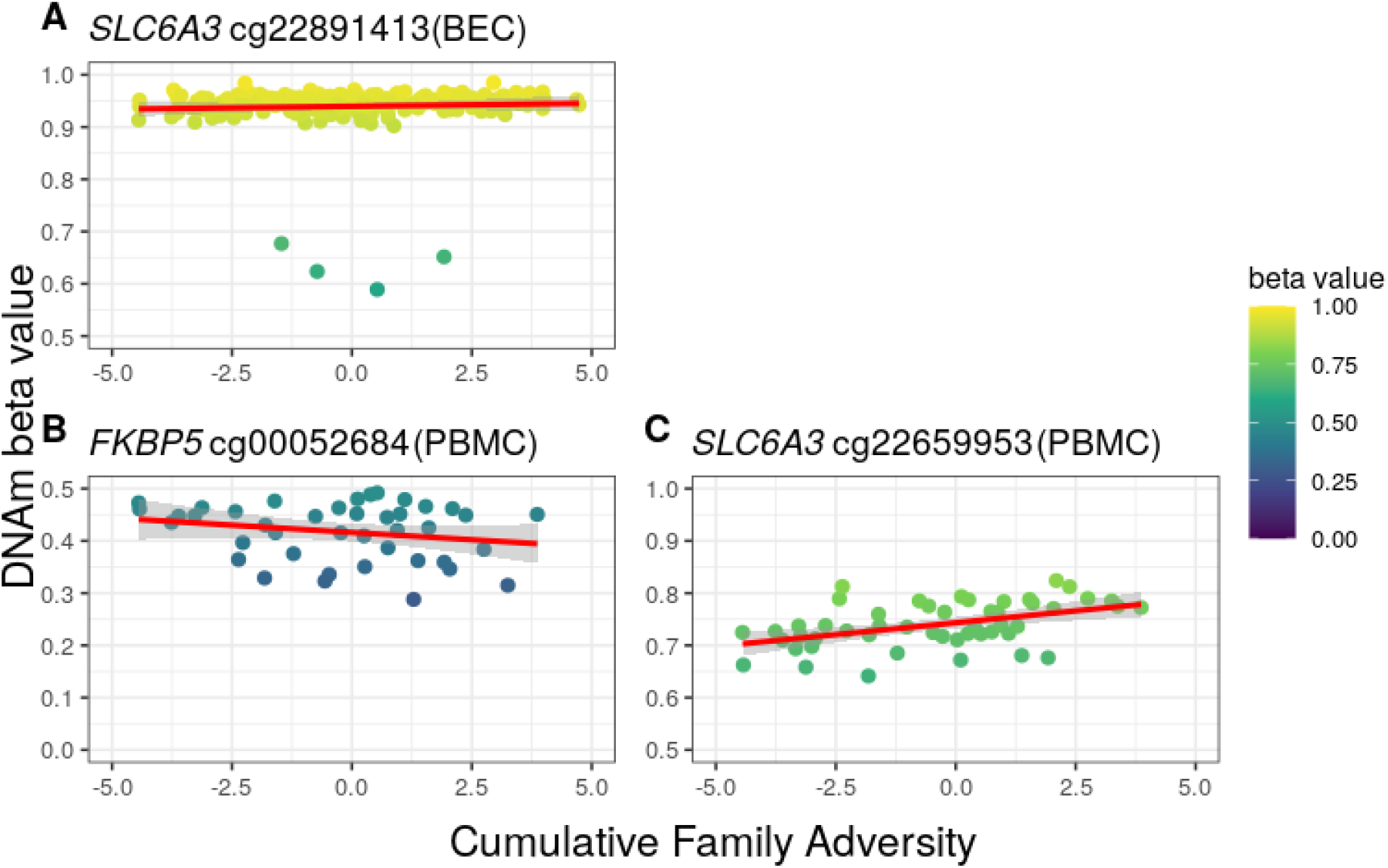
One and three candidate CpGs associated with cumulative family adversity (cumulative FA) passed the statistical threshold in BECs and PBMCs, respectively. Thirty candidate genes (366 and 323 variable probes in BECs and PBMCs, respectively) were identified for *a priori* analysis in both tissues. A regression model was run for each CpG to investigate the association between DNA methylation and cumulative FA while controlling for child’s sex, age, genetic ancestry, cell-type composition, and batch effect. Line graphs showing the relation between cumulative FA on the x-axis and β values on the y-axis for (A) the candidate CpG (cg22891314) in BECs that passed the statistical threshold (FDR<0.2) but not the technical threshold (|Δβ>.03|), and (B, C) two candidate CpGs (cg00052684 and cg22659953) in PBMCs that passed both the statistical and technical thresholds. The β values ranged from 0 to 1 with a y-axis scale of 0.5 to 1. The colors of the data points indicate β values. Trend lines and confidence intervals are shown.

**Table 3.** Seven Medium-Confidence DNA Methylation Sites in Children Associated with cumulative FA.

| CpG | *p* | FDR | *t* | $\Delta\beta$ | Chr | Gene | Genomic context | mQTL SNP | GeneMANIA association |
| --- | --- | --- | --- | --- | --- | --- | --- | --- | --- |
| cg22659953<sup>a</sup> | 1.78e-05 | 0.006 | 4.953 | .064 | 5 | *SLC6A3* | Gene body | No | 6 |
| cg00052684<sup>a</sup> | .001233 | 0.199 | -3.571 | -.066 | 6 | *FKBP5* | 5'UTR | No | 10 |
| cg04611670<sup>b</sup> | 4.73e-07 | 0.048 | 6.158 | .048 | 1 | *MEGF6* | Gene body | rs6697749 | 3 |
| cg17755321<sup>b</sup> | 1.89e-06 | 0.108 | 5.776 | .054 | 6 | *TNF* | 3'UTR | No | 13 |
| cg17487741<sup>b</sup> | 2.58e-06 | 0.108 | 5.552 | .040 | 13 | *RCBTB1* | 3'UTR | No | 4 |
| cg22664962<sup>b</sup> | 2.74e-06 | 0.108 | -5.557 | -.095 | 17 | -- | Intergenic | No | -- |
| cg23950278<sup>b</sup> | 3.17e-06 | 0.108 | -5.652 | -.069 | 11 | -- | Intergenic | No | -- |

<sup>a</sup> Candidate gene analysis. <sup>b</sup> Epigenome-wide association study.

### Five medium-confidence CpGs associated with cumulative FA discovered in exploratory site-by-site EWAS in PBMCs

Using 270,133 and 204,753 variable probes in BECs and PBMCs, respectively, we ran the same regression model as in candidate analyses (Tables S4, S5). After BH correction, no CpGs in BECs passed the statistical threshold of FDR<0.20 (*p*<2.00e−05) (Figure 2A). However, five medium-confidence CpGs in PBMCs showed significant associations with cumulative FA in our sample (Figure 2B, C)—cg04611670, cg17487741, cg17755321, cg22664962, and cg23950278—which were annotated to three genes: *MEGF6*, *RCBTB1*, and *TNF* (Table 3). One site, cg04611670, passed the statistical threshold of FDR<0.05 (*p*≤1.245e−7), operationally defined as a high-confidence site. In GeneMANIA, all 3 genes output from *MEGF6*, 3 of 4 output from *RCBTB1*, and 8 of 13 output from *TNF* were associated with immune function, while others were mostly related to neurological functions (Table S7).

**Figure 2.**
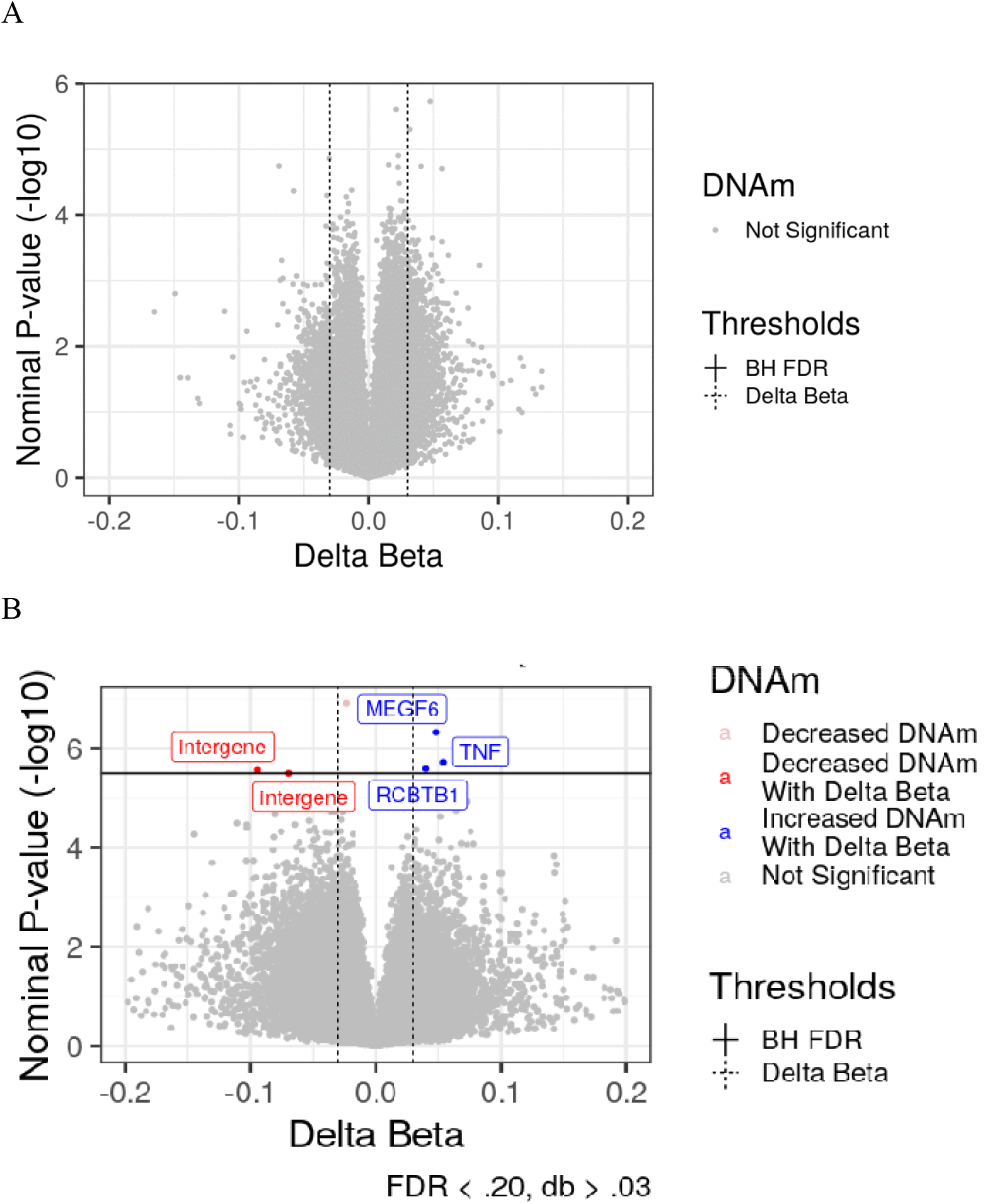
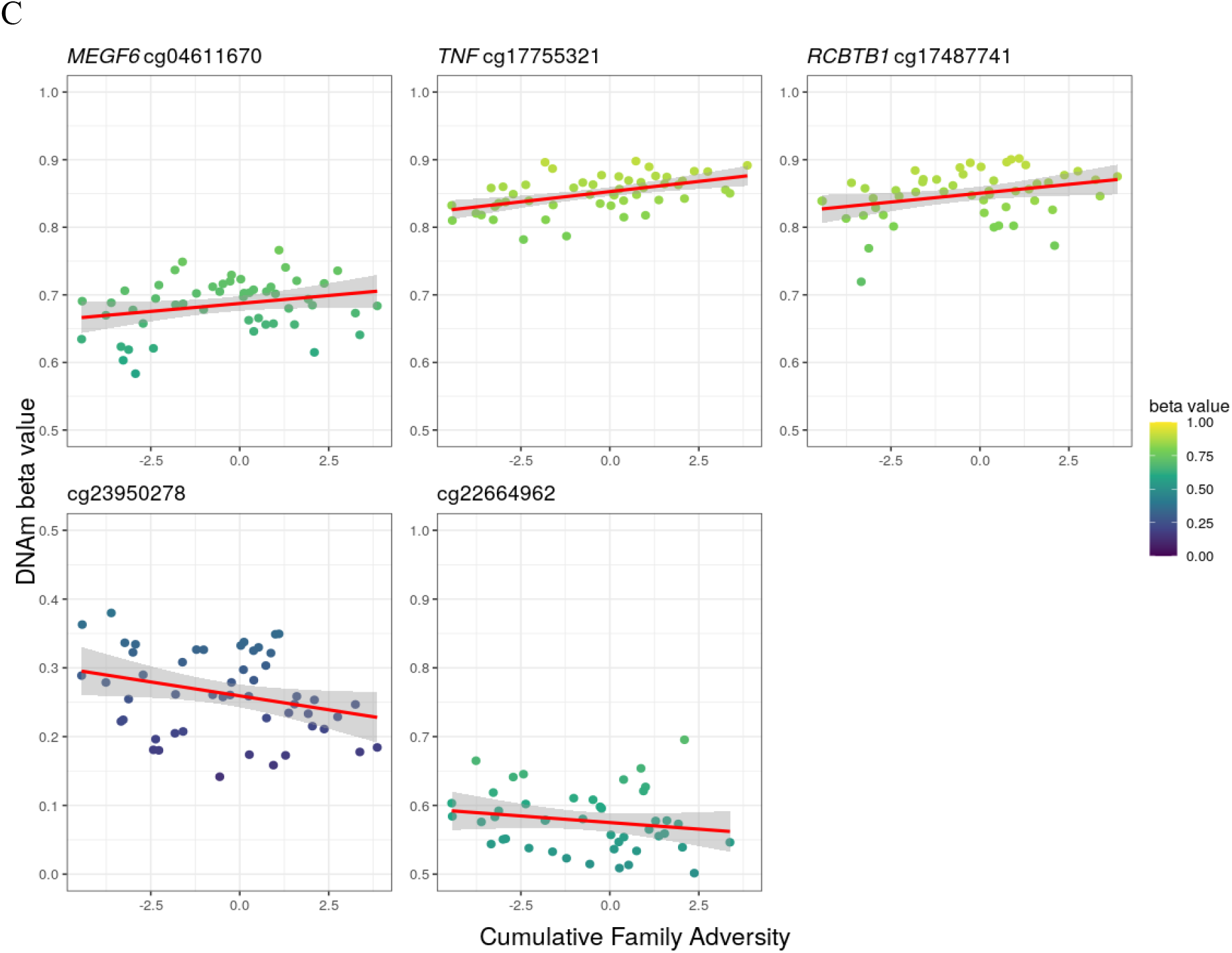
Results of site-by-site linear regression analysis of DNA methylation and cumulative FA (cumulative FA). The associations between DNAm level of each variable CpG site in BECs and PBMCs and cumulative FA were tested by linear regression while controlling for child’s sex, age, genetic ancestry, cell-type proportions, and batch effect, as depicted in the volcano plots. (A) Volcano plot showing that no CpGs in BECs passed the statistical (FDR<0.2) and technical thresholds (|Δβ>.03|), represented by the solid and dashed lines, respectively. (B) Volcano plot showing that five medium-confidence CpGs were significantly associated with cumulative FA, passing both statistical and technical thresholds. (C) Line graphs showing the relation between cumulative FA on the x-axis and β value on the y-axis for the five medium-confidence CpGs in PBMCs. The β values ranged from 0 to 1 with a variable y-axis scale of 0.5. The colors of the data points indicate β values. Trend lines and confidence intervals are shown.

### Tissue specificity of DNA methylation across BECs and PBMCs

Comparison of the overall Δβ distributions revealed greater differences in DNAm associated with cumulative FA in PBMCs (mean=−0.0035, SD=0.017, median=−0.0028) than BECs (mean=0.00068, SD=0.0091, median=0.000164). The same pattern was observed by comparing the 186,618 probes that were variable in both tissues (PBMC: mean=−0.0035, SD=0.017, median=−0.0028; BEC: (mean=−0.00075, SD=0.0097, median=0.00039). We investigated cross-tissue signals associated with cumulative FA using the matched tissue samples from 50 children. We examined the association between cumulative FA and DNAm at the seven medium-confidence CpGs identified in PBMCs in matched BEC samples. One of the seven sites (cg17755321) was invariable in BECs, and therefore cross-tissue concordance was not examined. Only one of the six remaining sites, cg17487741 annotated to *RCBTB1*, was significantly associated with cumulative FA with nominal *p*<.05 and |Δβ>.03| in both tissues (Table 4), but it should be noted that the FDR after BH correction was 0.226 (i.e., larger than our statistical threshold), suggesting tissue specificity of signals associated with cumulative FA.

**Table 4.** Association Between DNAm and Cumulative FA in Matched Tissue Samples From 50 Children at CpGs Identified in PBMCs Across All Samples.

| Site | Chr | Gene | Genomic context | PBMC $p$-value | $\Delta\beta$ | BEC $p$-value | $\Delta\beta$ |
| --- | --- | --- | --- | --- | --- | --- | --- |
| cg22659953 | 5 | *SLC6A3* | Body | 7.66e-04 | -.069 | .818 | .004 |
| cg00052684 | 6 | *FKBP5* | 5'UTR | 1.15e-06 | .049 | .743 | -.01 |
| cg04611670 | 1 | *MEGF6* | Body | 3.15e-05 | .040 | .104 | .037 |
| cg17755321 | 6 | *TNF* | 3'UTR | 3.88e-06 | .054 | -- | -- |
| cg17487741 | 13 | *RCBTB1* | 3'UTR | 2.36e-05 | .064 | .042 | -.06 |
| cg22664962 | 17 | -- | -- | 9.42e-06 | -.096 | .065 | .032 |
| cg23950278 | 11 | -- | -- | 4.29e-06 | -.071 | .878 | -.0023 |

We also examined cross-tissue concordance of DNAm levels regardless of their association with cumulative FA. We tested the correlations of DNAm levels across matched BECs and PBMCs at the medium-confidence CpGs found in PBMCs (Table 5). Significant positive correlations were found for only two of the six sites that were variable in both tissues, cg22659953 and cg22664962 (Figure 3), again indicating tissue specificity. The blood-buccal correlation provided by IMAGE-CpG [51] also showed that cg22659953 and cg22664962 had significant positive correlations between whole blood and buccal tissue. In addition, two other CpGs (cg04611670 and cg17487741) were reported to be significantly correlated in the database but were not significantly correlated in our sample (Supplementary Figure S4).

**Figure 3.**
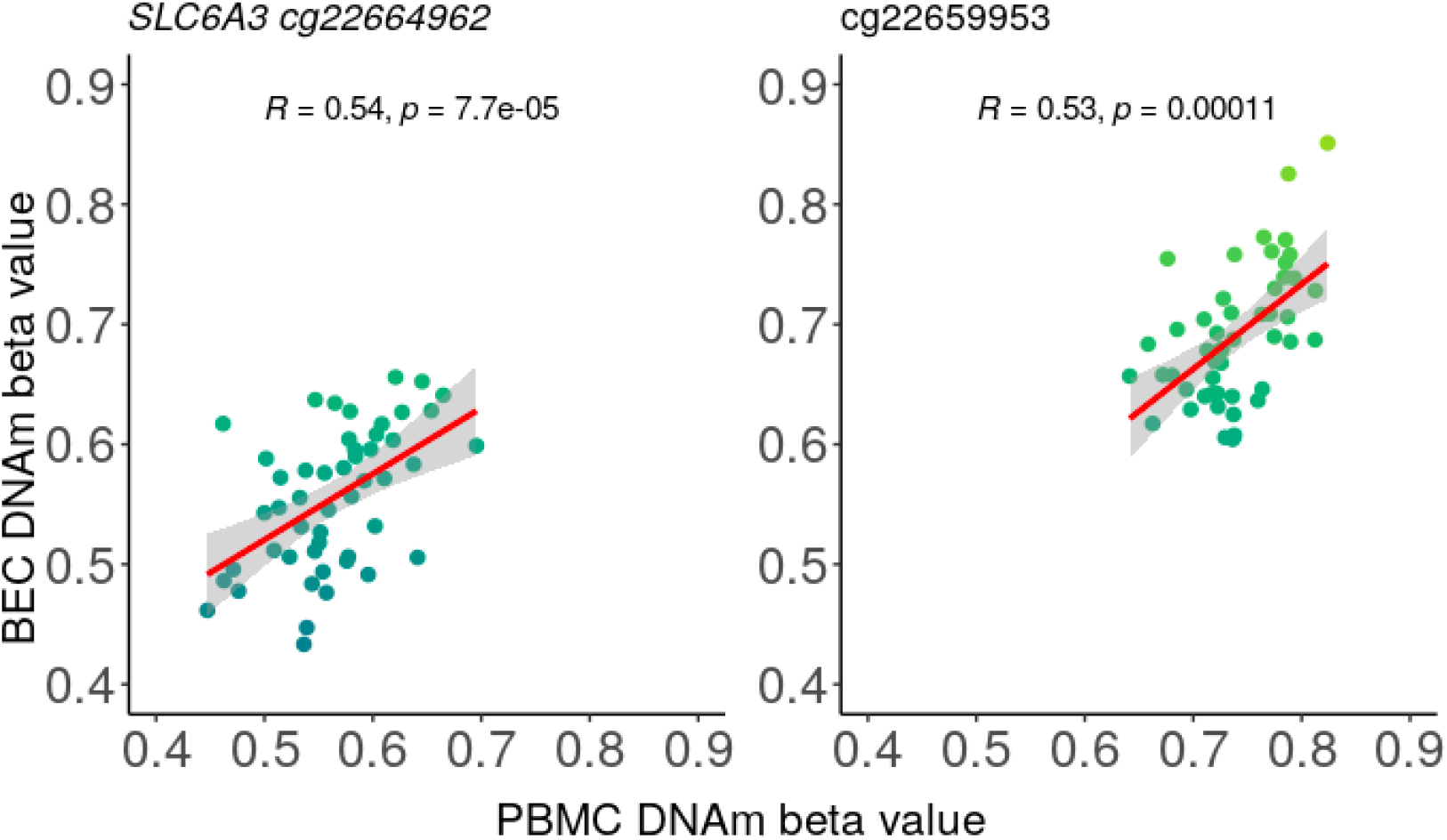
Scatter plot of DNAm at cg22659953 and cg22664962 in matched BEC and PBMC samples showing significant correlations.

**Table 5.** Correlation of DNAm Between Matched BECs and PBMCs in the Present Sample and IMAGE-CpG Database at Each CpG Found to be Associated With Cumulative FA.

| CpG | Mean $\beta$<br>BECs | Mean $\beta$<br>PBMCs | $\rho$ | $p$-value | IMAGE-CpG $\rho$ | IMAGE-CpG<br>$p$-value | Informative<br>CpG in<br>[45] |
| --- | --- | --- | --- | --- | --- | --- | --- |
| cg22664962 | 0.556 | 0.565 | 0.536 | 1.11e−4 | 0.662 | 0.003 | Yes |
| cg22659953 | 0.690 | 0.739 | 0.526 | 7.73e−5 | 0.651 | 0.002 | No |
| cg23950278 | 0.801 | 0.263 | 0.148 | 0.304 | −0.232 | 0.099 | -- |
| cg00052684 | 0.228 | 0.442 | −0.145 | 0.315 | -- | -- | -- |
| cg04611670 | 0.816 | 0.685 | −0.091 | 0.527 | 0.478 | 0.030 | No |
| cg17487741 | 0.895 | 0.848 | 0.041 | 0.775 | 0.690 | 0.001 | No |
| cg17755321 | 0.928 | 0.850 | -- | -- | -- | -- | No |

In a previous study scrutinizing cross-tissue concordance across BEC and PBMC samples in the same cohort [45], cg22664962 was also reported to be an informative site, defined as variable across individuals and highly correlated between BECs and PBMCs (i.e., 2 SD higher than the mean of the distribution; ρ>0.47), while cg22659953, cg04611670, cg17487741, and cg17755321 were reported to be differentially methylated across BECs and PBMCs, defined as FDR<0.05 and Δβ>.05.

It is also important to assess how these differential DNAm signals in blood, mostly relevant to stress and the immune system, may be reflected in the brain, which would have stronger implications for psychiatric outcomes [51,66]. As blood is a peripheral tissue and its DNAm level may not always be directly related to that in the brain, we examined the blood-brain DNAm concordance using BECon [66] and IMAGE-CpG [51] (Table S6, Figure S7). While two CpGs (cg00052684 and cg23950278) were not available in the BECon database, DNAm at cg22664962, cg17487741, and cg22659953 showed significant positive correlation with that in brain (mean correlations across all brain regions: *r*=0.40, 0.14, and 0.09, respectively). In contrast, DNAm at cg04611670 and cg17755321 in blood showed significant negative correlations with that in brain (mean correlations across all brain regions: *r*=−0.17 and −0.14, respectively). These correlations were all above the 50^th^ percentile of all CpG correlation values, suggesting that these CpGs were more strongly correlated in brain and blood than average; in particular, cg22664962 was in the 90^th^ percentile. In addition, all except two CpGs (cg22659953 and cg17755321) were defined as variable in the brain (>1% change in β value). Two CpGs (cg00052684 and cg17755321) were not available in the IMAGE-CpG database, which assessed different brain regions than BECon. DNAm at two CpGs (cg22659953 and cg22664962) showed significant positive correlations across brain and blood (Spearman’s correlation coefficients across all brain regions: ρ=0.456, *p*=.039 and ρ=0.555, *p*=.010, respectively).

### MEGF6-related SNP rs6697749 was an mQTL for cg04611670

The effects of the underlying genetic landscape on the CpGs found to be associated with cumulative FA were taken into consideration due to the well-documented genetic effects on DNAm through mQTLs [44,45]. A series of ANOVAs were run between the seven medium-confidence CpGs and all variable SNPs within a 5-kb window of their genomic location (Table S3). Only one CpG (cg04611670) was found to be significantly associated with a nearby SNP, rs6697749, related to *MEGF6* (F=7.25, FDR=0.10). The *post hoc* Tukey’s test showed that the β values were only significantly different across AA and BB allele genotypes (Δβ=.075) (Table 3, Figure S5). As there were only two samples with the AA genotype, further corroboration of this SNP as an mQTL was needed. A screen of the mQTL Database identified no *cis*-mQTLs within 10 kb of either cg04611670 or the other CpGs. Finally, cg04611670 did not have a bi- or trimodal distribution, suggesting that it was unlikely to be affected by mQTLs (Figure S6).

The potential mQTL SNP (rs6697749) was not significantly associated with cumulative FA in our PBMC samples (*F*(1,47)=0.307, *p*=.582) and did not contribute significantly to the association between cumulative FA and DNAm at cg04611670 (*p*=.848), as shown by comparing the effect size of the base model (0.00199) to that with rs6697749 (0.00233).

### Negative mother-child interaction contributed to relation of DNAm at SLC6A3-related medium-confidence CpG with cumulative FA

We examined the contribution of mother-child interaction quality to the association between children’s DNAm and cumulative FA at the seven medium-confidence CpGs identified in PBMCs. The two mother-child interaction quality PCs created from parenting variables available in this cohort were correlated with cumulative FA in the expected direction, with the negative interactions PC showing a positive correlation and the positive interactions PC showing a negative correlation with cumulative FA, although they did not meet the significance threshold (Pearson’s *r*=0.178, *p*=.226; *r*=−0.126, *p*=.391, respectively). Nevertheless, given the strong link between cumulative FA and mother-child interaction quality suggested in the literature [29,70], we examined the contributions of these interactions to the relation of DNAm and cumulative FA as a possible suppressor or amplifier [71]. Although no significant contribution was observed for positive mother-child interactions (Table S8), negative interactions showed a contribution of 21.69% to the association between cumulative FA and DNAm at cg2265995 (Table 6).

**Table 6.** Negative Mother-Child Interactions Contributed to the Relations Between One of Seven CpGs and Cumulative FA.

| CpG | Negative interactions<br>*p*-value | Negative interactions<br>FDR | cumulative FA effect<br>size | cumulative FA +<br>negative interactions<br>effect size | Negative interactions %<br>contribution |
| --- | --- | --- | --- | --- | --- |
| cg22659953 | 0.006 | 0.042 | 0.010 | 0.008 | 21.69 |
| cg00052684 | 0.315 | 0.441 | −0.010 | −0.011 | −6.19 |
| cg04611670 | 0.212 | 0.441 | 0.007 | 0.007 | 1.1 |
| cg17487741 | 0.280 | 0.441 | 0.006 | 0.006 | 3.13 |
| cg23950278 | 0.147 | 0.441 | −0.010 | −0.010 | 0.14 |
| cg17755321 | 0.460 | 0.537 | 0.008 | 0.009 | −14.05 |
| cg22664962 | 0.639 | 0.639 | −0.014 | −0.016 | −12.99 |

cumulative FA, cumulative family adversity.

## Discussion

Cumulative FA may get “under the skin” of children through biological pathways and influence their long-term health and development. Here, we examined DNAm in school-age children associated with cumulative FA in two common peripheral tissues, BECs and PBMCs, and found medium-confidence associations primarily in blood DNAm, suggesting strong tissue specificity. These connections were partially explained by negative mother-child interactions, suggesting the contribution of direct parent-child interaction quality to biological embedding of the family environment.

### Children’s blood DNAm in stress- and immune-related genes are associated with cumulative FA

To investigate the associations of DNAm with cumulative FA, we first examined stress-related candidate genes. Two medium-confidence DNAm CpGs associated with cumulative FA passed our statistical and technical thresholds in children’s PBMCs, but only one buccal DNAm CpG passed the statistical but not the technical threshold. One identified CpG was annotated to *FKBP5*, a stress-responsive gene and an important regulator of the hypothalamic-pituitary-adrenal (HPA) axis [72]. DNAm in *FKBP5* has been robustly linked with early-life adversity and stress-related psychiatric symptoms, such as posttraumatic stress disorder (PTSD) and other anxiety disorders [10,73–76], and *FKBP5* is also included in gene networks linked to GR signaling and inflammatory response [77]. One CpG identified in PBMCs and one CpG in BECs reside in *SLC6A3* encoding dopamine transporter, which has been implicated in a range of psychiatric disorders, including mood disorders, PTSD, schizophrenia, drug addiction, and oppositional defiant disorder in adults [78,79], as well as emotional and behavioral functioning in children [22,42,80,81]. DNAm in *SLC6A3* was also shown to be associated with other adverse environmental conditions, including child abuse and maternal smoking [82,83], and *SLC6A3* is part of a network of genes important for neurological functioning, including neurotransmitter regulation. In addition, the specific *SLC6A3*-annotated CpG in our study was linked to fetal alcohol spectrum disorder in two previous EWASs [84,85], suggesting its potential involvement in early development. Therefore, our candidate gene investigation replicated some previous DNAm findings related to early-life adversity and psychiatric symptoms, and provided support for our hypothesis of the biological embedding of cumulative FA through DNAm in stress-associated genes, potentially even across tissues in the case of *SLC6A3*.

Beyond the stress-related candidate CpGs, we discovered five medium-confidence DNAm CpGs linked to cumulative FA using an epigenome-wide approach in PBMCs. The changes in DNAm level at all these sites were small, especially in BECs. However, previous studies argued that small effects in human social epigenetic research can still be meaningful for children’s health and development [86]. Furthermore, the identified sites all passed a stringent technical threshold that was replicated in previous studies [62,63]. Several medium-confidence CpGs from the exploratory EWAS discovery identified in PBMCs are associated with immune function or reside within or near immune-related genes. Specifically, *TNF*, a key immune gene encoding tumor necrosis factor alpha (TNF-α) [87], has been linked to early-life adversity, such as childhood maltreatment [2,7,8]. Furthermore, differential DNAm of *TNF* was shown to be associated with early protective parenting and later proinflammatory responses [10,24]. In particular, the *TNF*-annotated CpG identified in the present study was also reported in a study of the relation between DNAm and mother-infant attachment style [63], corroborating its role in biological embedding of early social experience. Similarly, DNAm in *RCBTB1* encoding RCC1 and BTB domain-containing protein, a disease-related gene (e.g., various cancers) mainly expressed in basophils, was reported to be linked to negative environmental exposures, such as air pollution [88]. Both *RCBTB1* and *MEGF6* were also suggested to be involved in inflammation and immune-related gene networks, while one of the intergenic CpGs identified here (cg23950278) also resides close to *TRAF6* encoding TNF receptor-associated factor. In addition, differential DNAm of the two intergenic CpGs was implicated in early-life exposures, such as maternal folic acid supplementation and placental microbes [82,89]. Taken together, our findings indicated connections between early-life stress and DNAm signals in inflammation- and immune-related genes. As all signals were discovered in PBMCs, containing a high proportion of immune cells, it was unsurprising to find significant associations in immune-related genes.

Furthermore, we examined whether these DNAm differences can be explained by genetic effects. Although we found a potential mQTL (*MEGF6*-related SNP rs6697749) for one site (cg0461167), the differences in DNAm were only statistically significant across genotypes BB and AA, which represented a small subset of participants (genotype AA: *n*=2). Furthermore, this potential mQTL was not corroborated in mQTL Database or by the traditional bimodal or trimodal β distribution [68]. Therefore, caution is required when interpreting the potential genetic effects on DNAm at this site. Despite the well-known allelic effect on *FKBP5* methylation, the identified CpG annotated to *FKBP5* in our study was not significantly associated with any mQTL and was not reported in mQTL Database [90].

Our results were consistent with existing theoretical models and accumulating evidence of dysregulated stress and immune responses in relation to early cumulative FA. Early-life adversity, especially in the family environment, has long been linked to elevated risk of psychopathology [91–93]. One pathway was proposed to involve the disruption of response tendencies of the HPA axis, with daily stressful experiences at home leading to long-term dysregulated adaptations in the HPA axis, predicting mental health problems later in life [17,94]. Our findings added to the existing literature by suggesting that DNAm in genes involved in the HPA axis (e.g., *FKBP5*) may underlie these dysregulated adaptations, leading to increased susceptibility to psychopathology.

Furthermore, early-life adversity and dysregulated stress have been linked to poor physical health, especially through inflammation and immune pathways [7]. Environmental programming models, such as the Developmental Origins of Health and Disease (DOHaD) [95] and the Predictive-Adaptive Response model [96], posited that children may develop phenotypes considered adaptive to stimuli in their early environment but contribute to later adverse health outcomes. Specifically, exposure to cumulative FA in early life may induce a chronic proinflammatory state in children, reflected by changes in the proportion of immune cells in blood, increased cytokine levels, and DNAm of key immune-related genes [2,7,97]. Indeed, several studies showed that early-life stress exposure was associated with elevated levels of both CRP and IL-6 in children [98,99]. Importantly, a recent meta-analysis showed that the association between childhood stress and inflammation became stronger across the life course [7]. The present study provided further support for the contribution of early cumulative FA to the inflammatory state and immune health of children through differential DNAm at key immune-related genes, although further studies are required to replicate these results and investigate their potential functional significance.

### Negative mother-child interactions contribute to the association of children’s blood DNAm with cumulative FA

Given that daily experience of children may be more directly related to mother-child interaction quality, which can be negatively impacted by adverse family environments, the present study examined the roles of the quality of direct mother-child interactions on biological embedding of cumulative FA in children. Our results showed that negative mother-child interactions explained a significant portion of the associations of DNAm with cumulative FA, specifically at the medium-confidence CpG residing in *SLC6A3*, in both PBMCs and BECs, although the latter did not survive multiple-test correction. Therefore, negative mother-child interactions may serve as a behavioral pathway for the linkage of cumulative FA with DNAm patterns implicated in adult psychopathology and poorer emotional and behavioral functioning in children [100]. This aligns with the substantial evidence for the prevailing hypothesis regarding the connections between early adversity, negative parenting, and later psychopathology [2,29,101]. Nonetheless, the lack of significant correlations between negative mother-child interactions and cumulative FA in our sample was somewhat encouraging, as it suggested that mother-child interaction quality was not always determined by exposure to adversity. Indeed, the identification of negative mother-child interactions as a possible contributing factor illuminates potential avenues for future interventions, such as family therapy to support positive mother-child interactions and thus mitigate any potential negative link between cumulative FA and children’s biological patterns.

### Cross-tissue concordance across BECs, PBMCs, and brain

Using matched PBMCs and BECs, only one CpG was significantly associated with cumulative FA and showed similar changes in methylation level in both tissues, although this site did not survive FDR correction. As mentioned above, PBMCs represented the primary tissue in which DNAm patterns linked to cumulative FA were found in our study, despite the small sample size, which is often associated with insufficient statistical power to detect effects. Effect size is another major component of power calculations, and the power issue may have been alleviated because larger effect sizes were detectable in blood, as indicated by the greater overall Δβs in PBMCs than in BECs. These larger effect sizes in blood could be due to the greater cell-type heterogeneity in this tissue and its stronger relevance to immune and inflammatory processes associated with cumulative FA. Furthermore, given the link between stress and immune health, our findings suggested that DNAm in PBMCs, especially in genes involved in regulating immune function and inflammation, may readily reflect the biological embedding of early-life stress. The stronger signals in PBMCs were also consistent with previous studies suggesting that these cells may be more sensitive to sociodemographic factors than BECs [102]. Our findings highlighted the importance of choosing a tissue relevant to the variable of interest, and in the case of cumulative FA, blood may be the most suitable sample type for future investigations.

Despite limited concordance of DNAm with cumulative FA across tissues, further cross-tissue comparison showed that DNAm of more than half of the identified CpGs was significantly correlated across tissues with medium confidence according to IMAGE-CpG. Nonetheless, significant correlations of only two CpGs were also found in our matched samples, likely due to the small sample sizes. Furthermore, the blood-brain concordance database also indicated that most DNAm CpGs associated with cumulative FA in PBMCs were variable and significantly correlated with that in the brain. PBMCs were reported to share similarities in gene expression with the cerebellum across 4000 unique genes [103] and similar DNAm markers related to reward, stress responses, and psychopathologies, as identified in postmortem adult brain tissue [104]. Therefore, PBMC DNAm patterns associated with cumulative FA may also be reflected in brain DNAm, which could contribute to the later development of mental health problems. Further studies are required to determine the relation of DNAm in the developing brain to that in blood as a more readily accessible peripheral surrogate tissue.

### Limitations

This study had several limitations that should be taken into account when interpreting our findings. First, the current study was conducted with children who lived in Canada, were primarily White, and had well-educated mothers. The experience of cumulative FA and how it may be associated with children’s development can be highly context-specific, so the present findings may not be generalizable to children from different and more diverse racial and socioeconomic backgrounds. Second, we adopted a moderately lenient threshold for multiple-test correction across all analyses (FDR<0.20), thus accepting some false positives. This threshold was implemented along with a stringent, robust technical threshold of |Δβ>.03| [63], adding confidence to our findings, as the difference observed was greater than technical noise likely resulting from the array methodologies used in the analyses. Nonetheless, given the small sample size and the thresholds used, further studies are required to corroborate these findings. Second, all data were obtained at the same time point, and therefore the associations between cumulative FA and DNAm were cross-sectional and correlational in nature. Although we hypothesized that the DNAm patterns may have been induced by cumulative FA, it is possible that they co-occurred due to other factors or that these DNAm patterns in children existed prior to exposure to stress. Therefore, these associations should be investigated longitudinally in future studies to scrutinize their temporal order and relations over time. Third, despite collecting samples from the same individuals, it should be noted that PBMCs were collected at a slightly later time point than BECs. As age and development are important drivers of DNAm differences, we controlled for children’s age in both models. However, the cross-tissue similarities could still be attenuated, and further studies with matched samples collected at the same time point are needed to confirm our findings. Finally, our cumulative FA construct was only derived from maternal reports, and therefore did not consider the paternal or child’s perspective regarding the social environment. Future studies adopting a more family system perspective with collection of data from multiple family members to gain a comprehensive understanding of the family environment will be beneficial to understand the relations of DNAm with biological embedding of cumulative FA.

## Conclusions

Despite the limitations outlined above, our study contributed to the existing literature regarding the associations of DNAm with cumulative FA by comparing the patterns in two common tissues in school-age children. This cross-tissue comparison suggested that PBMCs may be a relevant tissue type for identifying DNAm markers of early psychosocial stress. Extending previous studies on cumulative FA and DNAm, our robust analyses with consideration of important biological factors driving DNAm, including cell-type heterogeneity, genetic ancestry, and effects of genetic variants [11], supported the link of such exposure with DNAm, especially in stress- and immune-related genes. Further, we showed that direct parenting factors, such as mother-child interaction quality, represented possible pathways underlying this relation as potential avenues for future interventions.

## Summary points

- Cumulative family adversity in early life was shown to be associated with later mental and physical health outcomes.
- Cross-tissue comparison of pediatric DNA methylation patterns associated with early-life stress is limited, but important for guiding future investigations.
- The potential roles of proximal factors through which cumulative family adversity becomes biologically embedded are unclear.
- This study examined epigenome-wide DNA methylation patterns in 218 buccal samples and 51 blood samples linked to cumulative family adversity in school-age children.
- Differential DNA methylation associated with cumulative family adversity was only discovered in blood but not buccal tissue, with limited cross-tissue concordance and genetic influences.
- Negative mother-child interactions explained a significant portion of the association between blood DNA methylation and cumulative family adversity.
- Blood DNA methylation associated with cumulative family adversity was primarily found in genes involved in stress regulation and immune function.
- The findings supported DNA methylation in blood as a potential biomarker linking early psychosocial stress and later mental and physical health outcomes.

## Supporting information

Supplementary Material on DNA methylation data

