## Supplementary Material on DNA methylation data for "Cross-Tissue Specificity of Pediatric DNA Methylation Associated with Cumulative Family Adversity"

***DNA sample collection, extraction, and microarray genome-wide DNA methylation profiling***

All procedures were approved by the institutional review board at the University of British Columbia and were described previously [50]. Briefly, a parent or legal guardian of the child provided written informed consent and each child also provided verbal assent before inclusion in the study. BEC samples were obtained from children using Isohelix Buccal Swabs (Cell Projects Ltd., Kent, UK) and stabilized at room temperature with Isohelix Dri-Capsules prior to DNA extraction. Blood samples were collected at a follow-up visit ranging from 8 months to 2.06 years after buccal sample collection. PBMCs were isolated from whole-blood samples collected into Vacutainer® CPT™ Cell Preparation Tubes (Becton, Dickinson and Company, Franklin Lakes, NJ, USA), as described previously [50]. PBMC pellets were frozen and stored at −80°C prior to DNA extraction.

DNA was extracted from both buccal and PBMC samples and subjected to bisulfite conversion using an EZ DNA Methylation Kit (Zymo Research, Irvine, CA, USA); full details were described previously (Islam et al., 2019). The samples were then applied to the Illumina Infinium HumanMethylation450K Beadchip (450K) DNAm array in accordance with the manufacturer’s protocols (Illumina, San Diego, CA, USA), producing 485,577 data points exported as raw intensity IDAT files used for DNAm preprocessing.

***Microarray genome-wide SNP profiling***

Saliva samples were also collected from children to extract DNA for genome-wide single nucleotide polymorphism (SNP) profiling using the Oragene OG-500 DNA all-in-one system in accordance with the manufacturer’s protocol (DNA Genotek, Kanata, ON, Canada). Extracted DNA was applied to the Illumina Infinium PsychChip BeadChip genotyping array (PsychChip) with measurement of 588,454 SNP probes.

***Data processing***

**DNA methylation data preprocessing.** All tissue samples (376 BEC and 92 PBMC samples, in addition to 55 dried blood spots that were not used in the present study) were preprocessed together to reduce the impact of batch effects. The raw intensity values from the DNAm array were subjected to initial quality control checks for array staining, extension and bisulfite conversion, color correction, and background adjustment using Illumina GenomeStudio V2011.1 software. DNAm data were then exported as methylation beta values ranging from 0 (unmethylated) to 1 (methylated) for each CpG measured on the array, which were preprocessed using R version 4.0.5 and Bioconductor version 3.12.0. Quality control checks were completed using the ewastool and minfipackages [62,63]. Samples that failed two or more control metrics in ewastool, or failed both quality checks in ewastooland minfi were removed (*n* = 28). Additional checks to confirm the sex of the participants were performed using XY and SNP probes; two samples that had mismatched/uncharacterized sex prediction were filtered out. Further checks were conducted to remove samples with poor performance (*n* = 4) that failed two or more of the following checks: bead counts fewer than three beads/reads, detection *p* > 0.01, failed sample outlier detection, probability SNP outlier > 20%, and failed quality control metrics in ewastool or minfi. A total of 34 samples were removed after all quality control checks. To confirm that matched tissue samples were collected from the same individual, we used 65 SNP probes included on the 450K array to examine genetic relatedness. These probes were then removed from the data set. To account for type I and type II probe differences on the 450K array, normalization was performed using the minfi package preprocess *Funnorm* function [64]. Additional probe filtering was conducted on the normalized DNAm data, including probes predicted to cross-hybridize, probes that bind to the sex chromosomes, and probes with detection *p* > 10 as well as those with NAs in more than 2% of samples [65]. After all preprocessing steps and removal of duplicate samples from the same individuals (*n* = 10), 298 buccal and 81 PBMC samples, of which genotype data were available for 232 and 55, respectively, and 485,577 probes in each sample remained. Among these samples, only one child per family was selected for subsequent analysis with priority given to children who provided both buccal and PBMC samples. Overall, 218 buccal and 51 PBMC samples (all with genotype data), of which 50 were matched from the same individuals, and 485,577 probes in each sample were used for subsequent analyses.

**DNA methylation data reduction.** To reduce the testing space and address the equal significance likelihood assumption of the Benjamini–Hochberg (BH) False Discovery Rate (FDR) control method [66,67], we subset the DNAm probes using an interquartile range filter to include only variable probes where the methylation β value varied by at least 5% across samples in the 5^th^ and 95^th^ percentiles. This resulted in 270,133 and 204,753 variable probes that were included in subsequent analyses for BECs and PBMCs, respectively.

**Genotyping data preprocessing.** The raw genotyping data from the array were processed using the Genotyping Module of Illumina GenomeStudio software (version 2.0.4). Samples with more than 5% of genotyped SNPs missing were removed. SNP probe filtering was performed based on the Illumina guide for processing Infinium genotyping data. Briefly, SNP probes located on mitochondrial DNA, on sex chromosomes, or without chromosome labels were removed. SNPs will a call frequency < 97%, cluster separation < 45%, and minor allele frequency < 5% were removed. Finally, probes were filtered based on the Het Excess score, a measure of heterozygosity relative to the Hardy–Weinberg Equilibrium; probes with Het Excess score > 0.2 and < −0.3 were removed. After probe filtering, the SNP probe count was 262,327. From these probes, non-ACTG SNPs and duplicates were further removed, leaving 257,202 probes as the final count.

The 450K DNA methylation array also contains 65 SNPs that can be used for quality control. Genotyped samples from the PsychChip array were matched with samples measured on the 450K array by comparing allele frequencies of the 15 common SNP probes present on both arrays. To ensure samples were matched between the arrays, only those with > 90% correlation in allele frequencies were retained. All data filtration and preprocessing steps were performed in R version 4.0.5 and Bioconductor version 3.12.0.
