## Supplementary Figure S1-S7 for "Cross-Tissue Specificity of Pediatric DNA Methylation Associated with Cumulative Family Adversity"

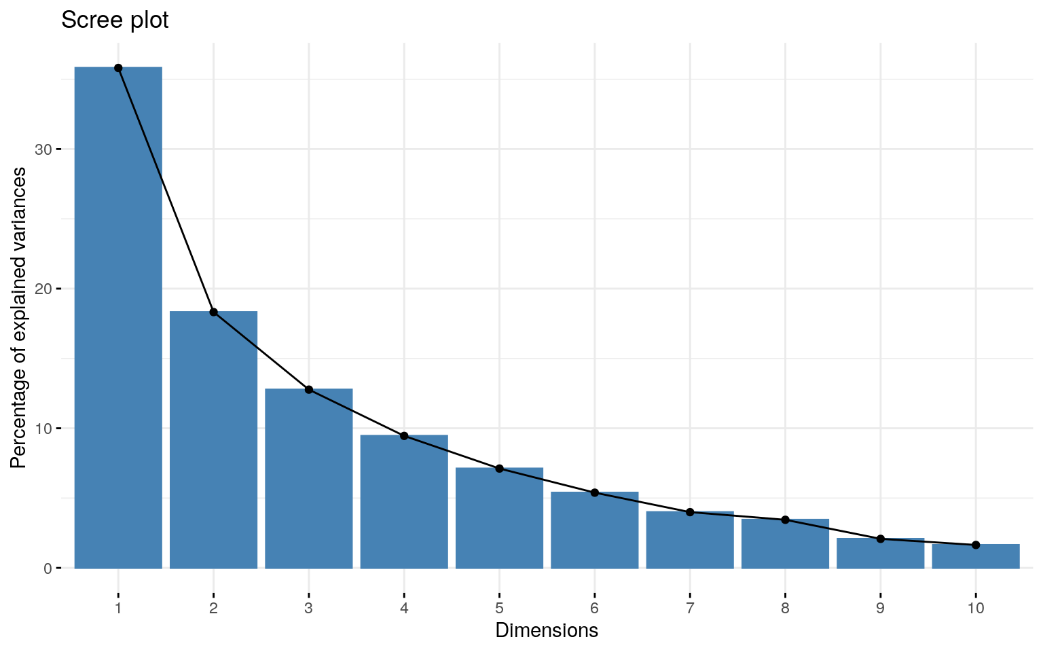

Figure S1. Scree plot suggesting a 2-factor solution

a
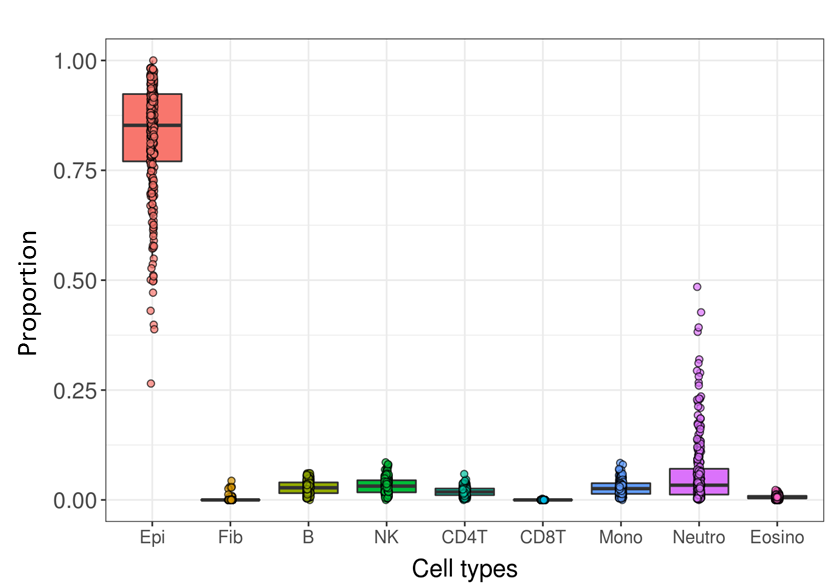

b
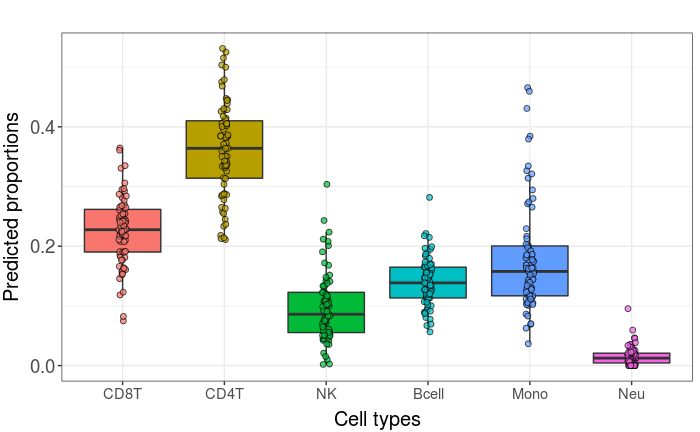

c
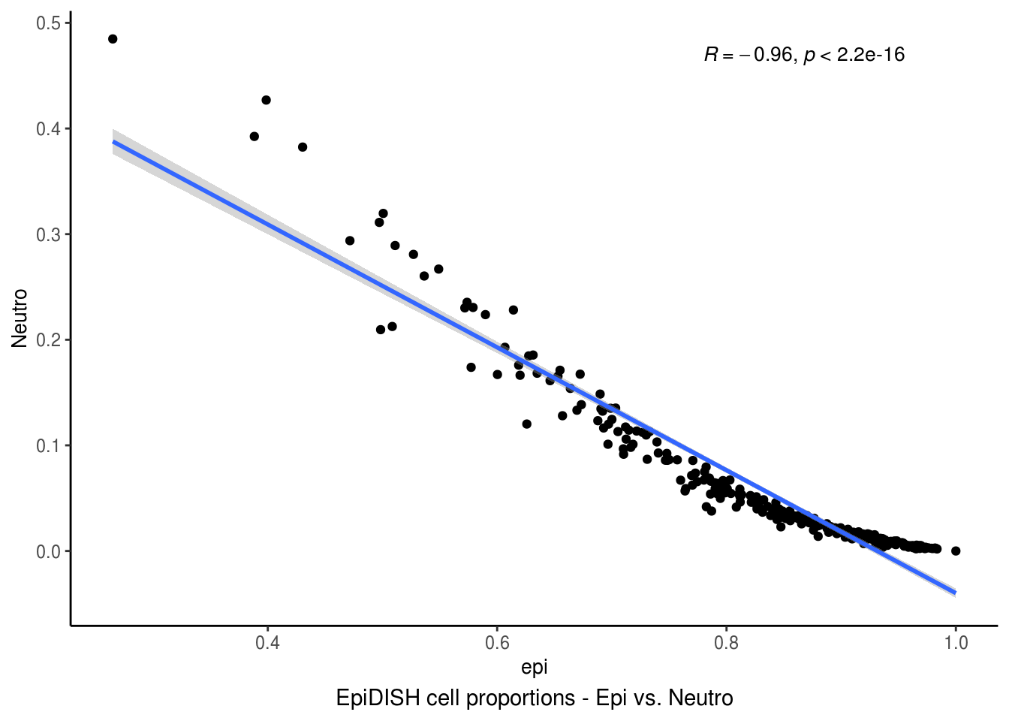

Figure S2. Cell type estimation from DNA methylation (a) Boxplot showing proportion of nine cell types estimated by HepiDISH in Buccal samples (b) Boxplot showing proportion of nine cell types estimated by Houseman IDOL in Blood samples (c) Scatterplot showing significant negative correlation between with epithelial cell and neutrophil proportions

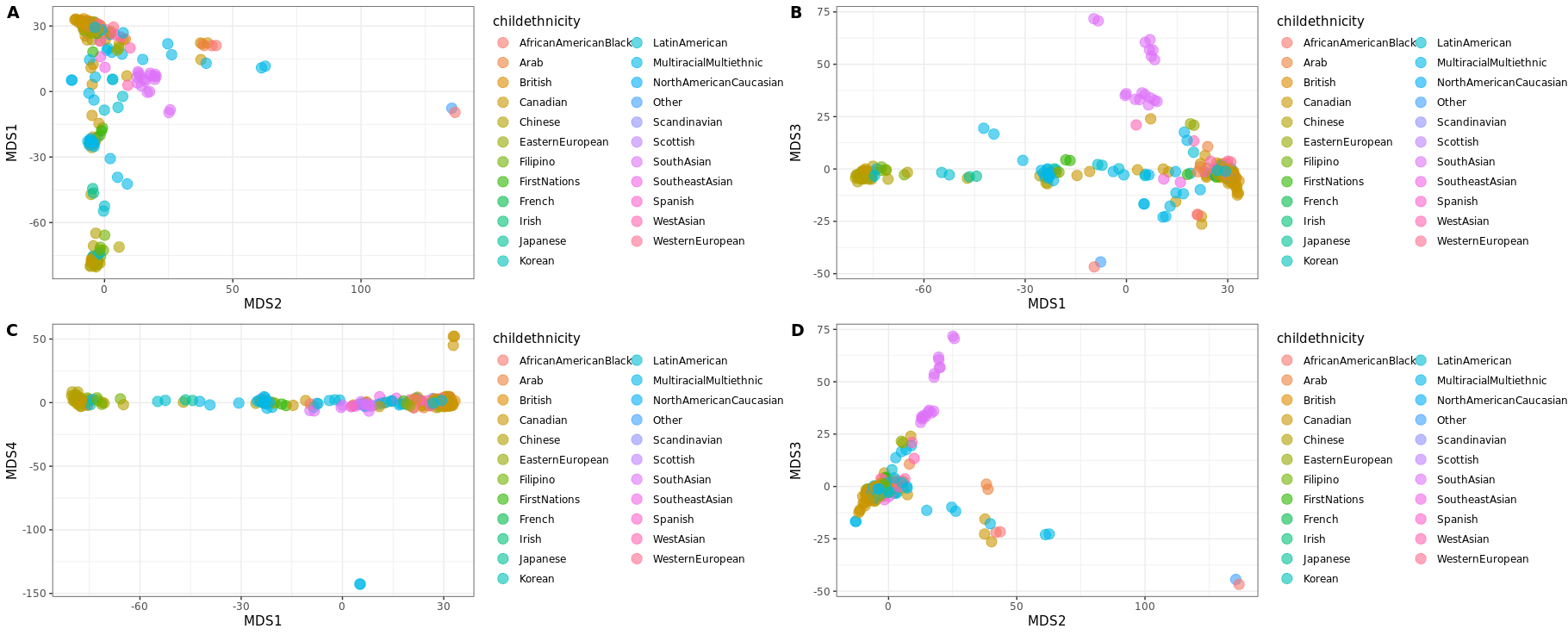

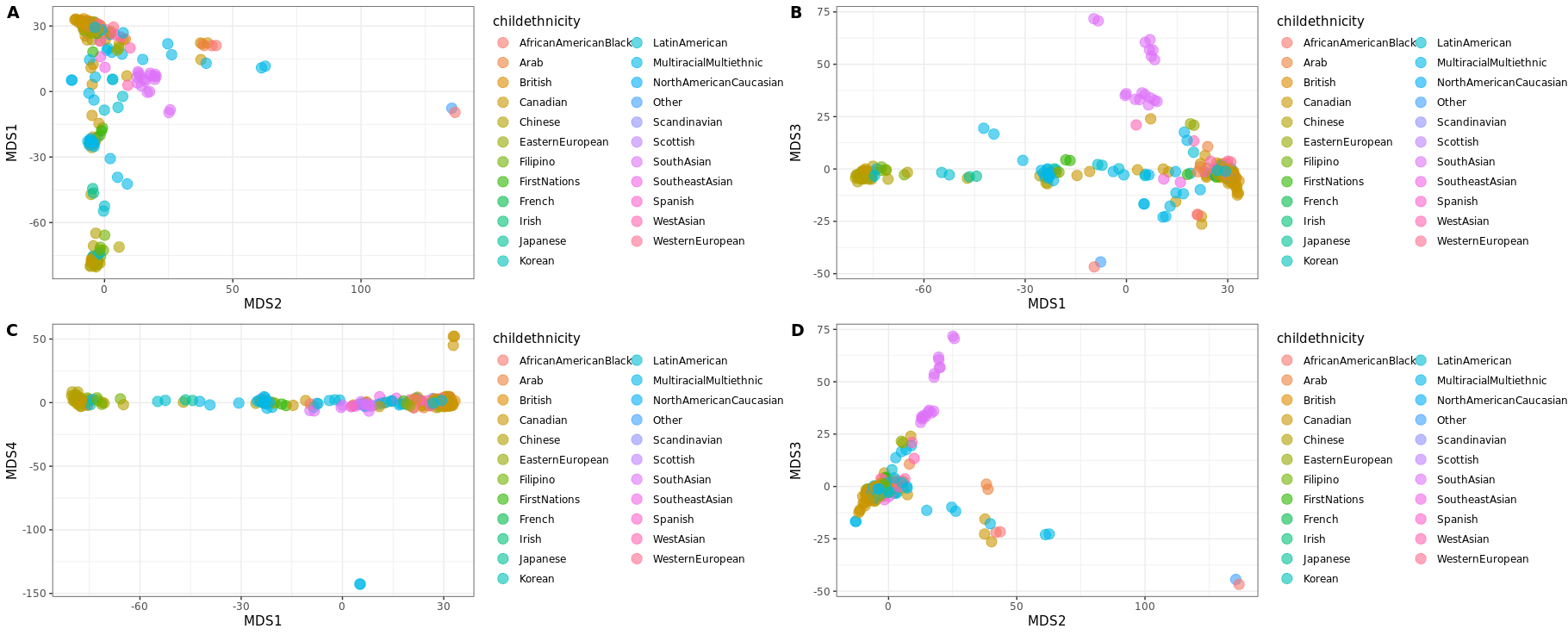

Parental reported child ethnicity

Figure S3. Plots of MDS coordinates with parental reported child ethnicity variables

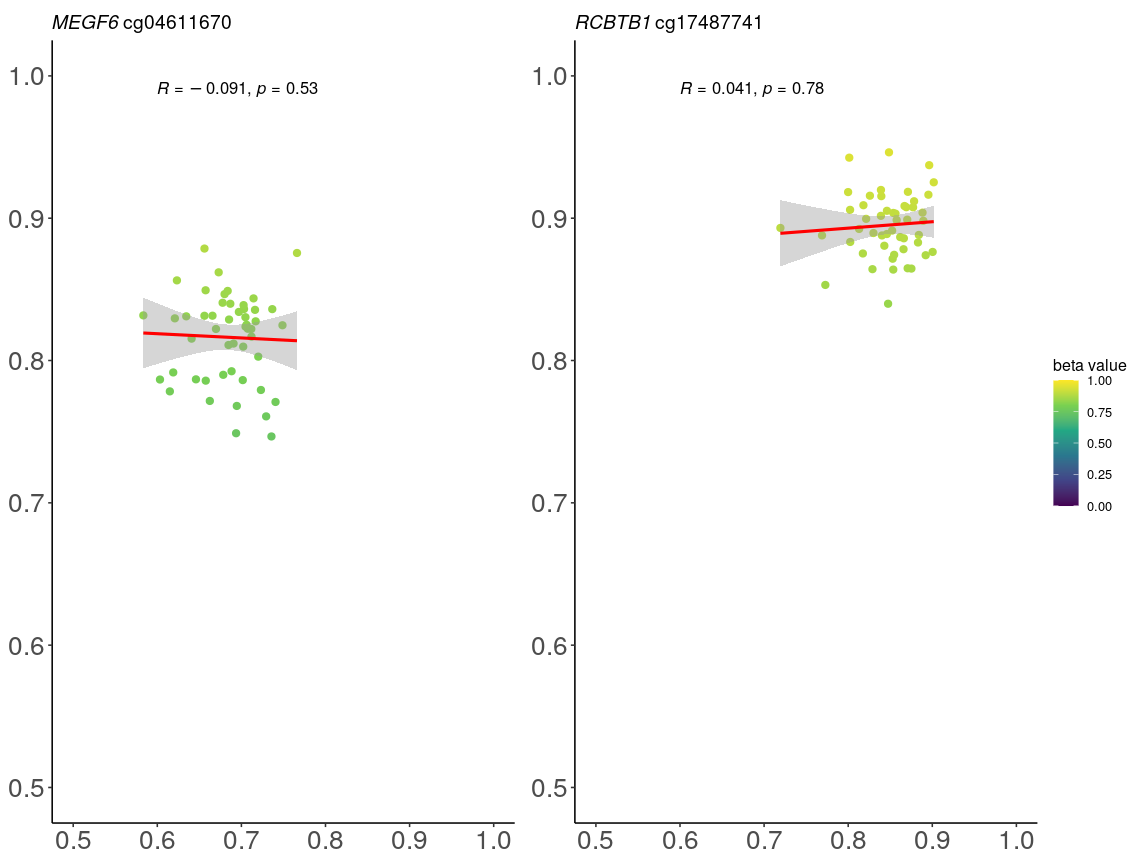

Figure S4. Scatterplot of BEC DNAm with PBMC DNAm in matched samples at cg04611670 and cg17487741. The correlation was not significant in our sample but significant according to IMAGE-CpG

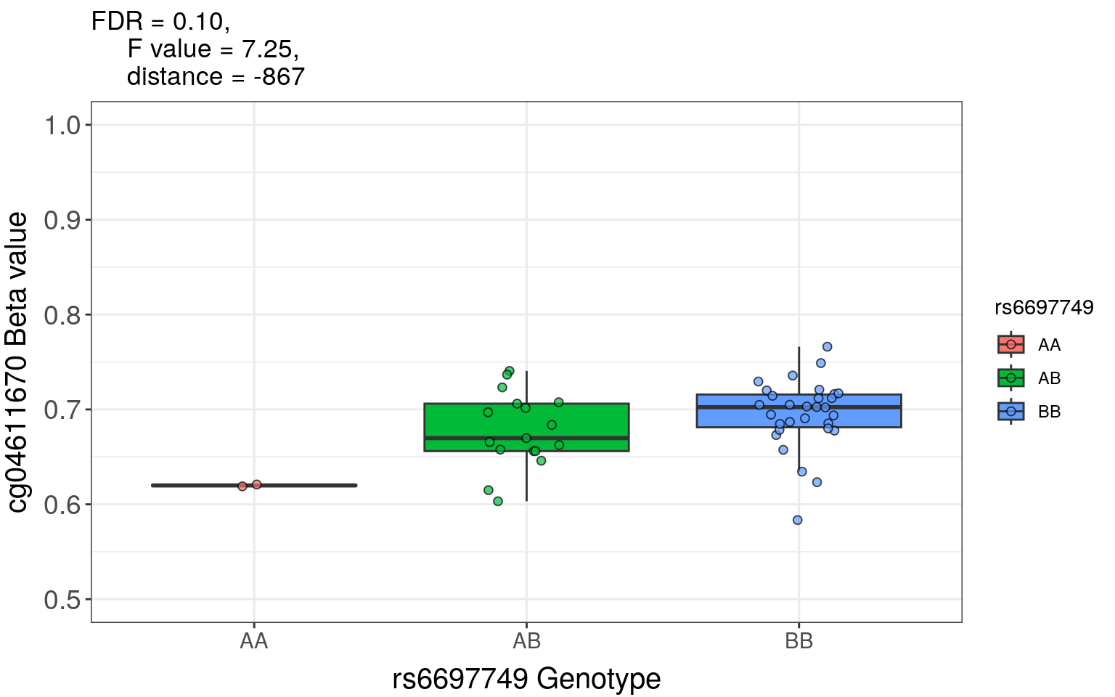

Figure S5. Boxplot showing beta value of cg04611670 differ across genotype of rs6697749

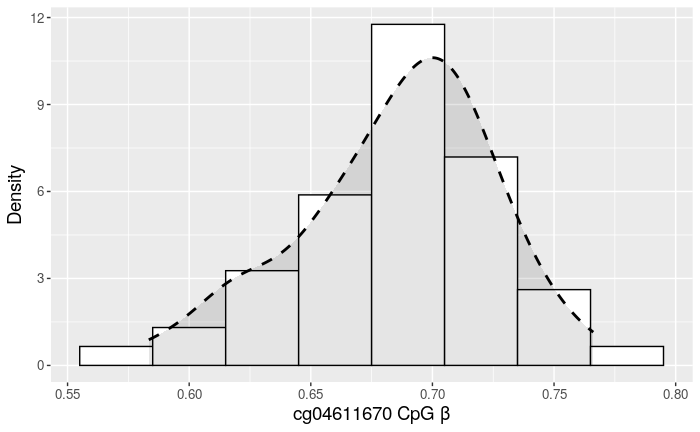

Figure S6. Beta distribution of cg04611670, which did not show a bi- or trimodal distribution

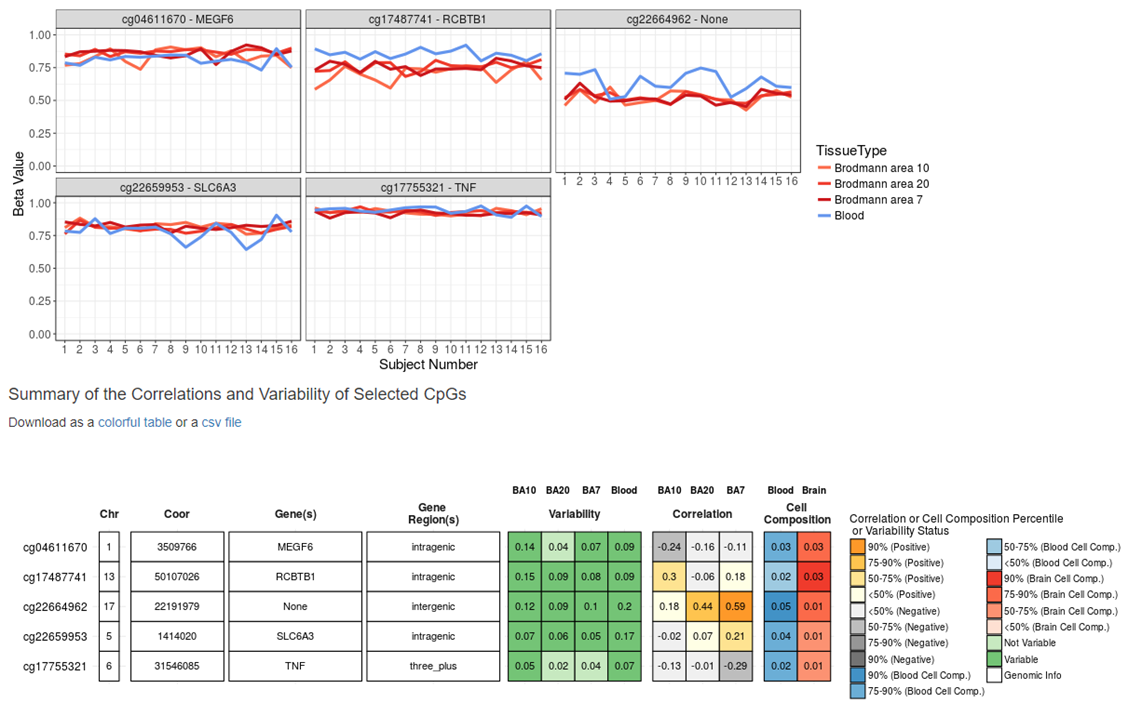

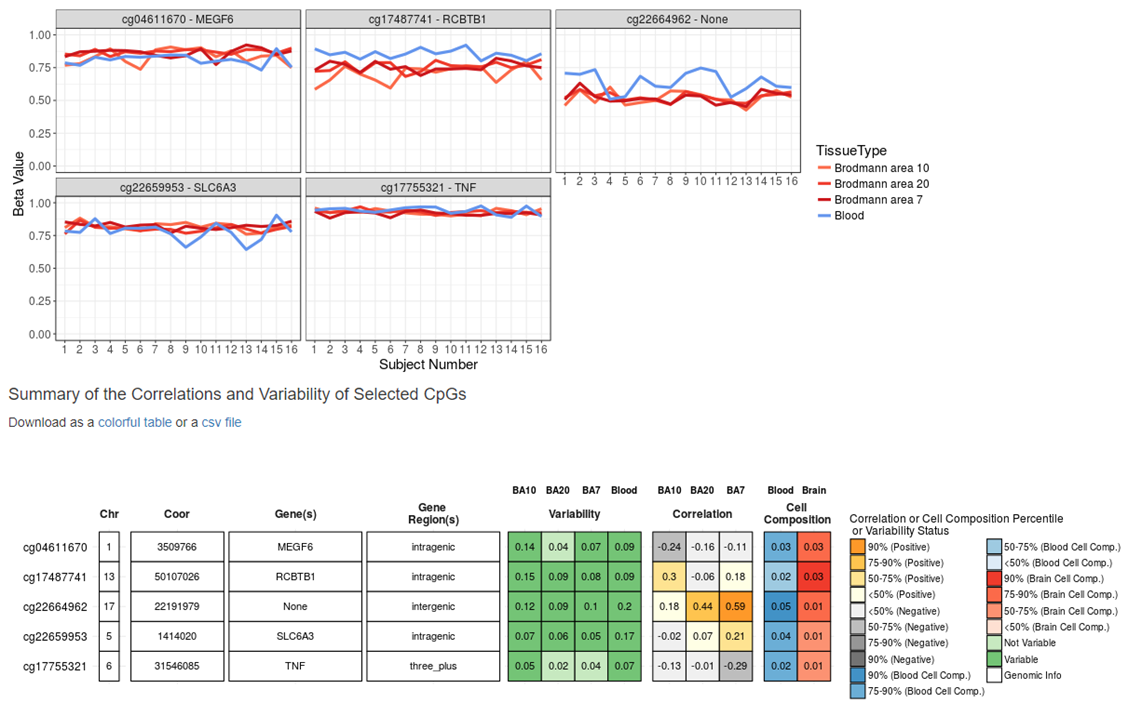

Supplementary material Figure S7. Blood and brain correlation for the five CpGs that were in the Blood-Brain Epigenetic Concordance (BECon) database
