## Supplementary Table 8 for "Cross-Tissue Specificity of Pediatric DNA Methylation Associated with Cumulative Family Adversity"

Supplementary Table S8. Positive Mother–Child Interactions did not Contribute Significantly to the Relations Between CpGs and Family Adversity

| CpG | Positive interactions *p*-value | Positive interactions FDR | Fam adv Effect size | Fam adv + interactions Effect size | Positive interactions % contribution |
| --- | --- | --- | --- | --- | --- |
| **cg22659953** | **0.042** | **0.296** | **0.010** | **0.008** | **17.95** |
| cg22664962 | 0.183 | 0.641 | −0.014 | −0.015 | −6.36 |
| cg00052684 | 1.000 | 1.000 | −0.010 | −0.011 | −8.92 |
| cg04611670 | 0.906 | 1.000 | 0.007 | 0.008 | −4.17 |
| cg17487741 | 0.866 | 1.000 | 0.006 | 0.006 | 1.97 |
| cg17755321 | 0.460 | 1.000 | 0.008 | 0.008 | −2.36 |
| cg23950278 | 0.789 | 1.000 | −0.010 | −0.011 | −7.51 |

Fam adv, cumulative family adversity.
